# Lymphangiogenesis driven by VEGF-C reshapes tumor immune landscape and enables tumor eradication by viral immunotherapy

**DOI:** 10.1101/2025.08.27.672466

**Authors:** Ruben Fernandez-Rodriguez, Sara Cuadrado-Castano, Andrew Edwards, Anita Rogic, Yonina Bykov, Ignacio Mena, Noelia Dasilva-Freire, Alice O Kamphorst, Rosa Karlic, Amir Horowitz, Adolfo García-Sastre, Mihaela Skobe

**Author notes:** These authors contributed equally to this work (Ruben Fernandez-Rodriguez, Sara Cuadrado-Castano).

## Abstract

Vascular Endothelial Growth Factor C (VEGF-C) is a key lymphangiogenic growth factor expressed by many types of cancer. Here, we investigated the effects of tumor-derived VEGF-C on the therapeutic efficacy of oncolytic virotherapy with avian paramyxoviruses APMV-1 (NDV) and AMPV-4. Treatment of B16F10 tumors not expressing VEGF-C with either virus led to tumor growth delay, and complete responses were rare. In contrast, when tumors expressed VEGF-C, viral therapy led to complete remission and long-term immunological memory in most mice. Upon re-challenge, most mice remained tumor- and metastasis-free for over 18 months, indicating durable immunity. VEGF-C induced tumor lymphangiogenesis, which correlated with high CD8^+^ and CD4^+^ T-cell densities in proximity of lymphatic vessels. Spectral flow cytometry revealed distinct changes in the composition and activation of CD8^+^, CD4^+^ T cells and NK cells associated with complete remission. In responders, tumors were highly enriched in CD8^+^CD25^+^ PD-1^+^ effector T cells. Composition of T cells was altered in sentinel and in contralateral lymph nodes, indicating a systemic immune response. Taken together, these data demonstrate VEGF-C-induced changes in tumor immune landscape which are critical for achieving complete response and support combining VEGF-C with APMV virotherapy as a novel highly effective strategy for cancer treatment.

## INTRODUCTION

Viral immunotherapy is an emerging cancer treatment that leverages the ability of certain viruses to break immune tolerance and stimulate anti-tumor immunity (*1*). A major challenge in clinical development of oncolytic viruses (OVs) has been their limited therapeutic efficacy as monotherapy, often leading to partial responses (*2*). Consequently, OVs are mainly employed in combination with immune checkpoint inhibitors, chemotherapy, or radiotherapy, to enhance therapeutic efficacy (*3*), and various strategies have been explored to enhance immune activation by OVs that directly target immune cells (*4–7*). To date, four OVs have been approved worldwide for cancer treatment (*1, 8–11*). These promising results in the clinic highlight the therapeutic potential of OVs as well as the need for novel strategies that improve efficacy while minimizing toxicity.

Avian avulaviruses, referred to as avian paramyxoviruses (APMVs) are negative-sense single-stranded RNA (ssRNA (-)) viruses (*12*) with oncolytic activity, strong immunostimulatory properties and an excellent safety profile (*6, 13–15*). Among them, Newcastle disease virus (NDV; APMV-1) has been most extensively studied and has garnered significant interest for cancer therapy (*14, 16–21*). APMVs preferentially replicate in tumor cells, exploiting their defective antiviral defenses (*1*). Their replication is restricted to the cytoplasm of the host cell, without DNA intermediates (*12*), which minimizes the risk of genomic integration and enhances safety. Importantly, APMVs are not human pathogens (*22*). Given their strong safety profile, inherent selectivity for cancer cells, and ability to elicit potent anti-tumor immunity APMVs demonstrate strong potential as cancer therapeutics.

Enhancing the anti-tumor efficacy of OVs requires a deeper understanding of the key determinants of immune response within the tumor microenvironment (TME), and much attention has been given to the interactions between tumor cells and various immune cell subsets. Tumor vasculature also plays an important role in shaping TME, and recently lymphatic vessels have emerged as modulators of immunity (*23–25*). Beyond serving as conduits for immune cell trafficking to the lymph nodes, they actively shape immune responses through chemokine production, scavenger functions, and antigen presentation among others (*26*). Lymphatics can promote both tolerance and immunity, and their impact is highly context dependent. Vascular Endothelial Growth Factor C (VEGF-C) is a key lymphangiogenesis factor (*27–29*), which activates lymphatic endothelial cells (LECs) through the tyrosine kinase receptors VEGFR-3 (*27, 30–33*) and VEGFR-2 (*27*). While initially preclinical studies demonstrated that VEGF-C activation of VEGFR-3 promotes an immunosuppressive microenvironment in tumors, more recent studies indicate that VEGF-C potentiates immunotherapy response (*34, 35*). These seemingly contradictory outcomes underscore the context-dependent influence of VEGF-C and lymphatics on tumor immunity.

Here, we investigate the effects of VEGF-C on therapeutic efficacy of APMVs in mouse melanoma. We show that VEGF-C markedly enhances anti-tumor immune responses induced by APMVs. Characterization of tumor immune landscape demonstrates unique changes of immune cell subsets in B16F10 tumors expressing VEGF-C and treated with APMV-1 (NDV), that were associated with complete response. This study demonstrates that VEGF-C potently enhances the effects of OVs leading to durable anti-tumor response and immunological memory.

## RESULTS

### Oncolytic viral therapy with APMVs leads to complete remission of B16F10 tumors expressing VEGF-C

We used NDV and APMV-4 to examine their effectiveness against tumors expressing VEGF-C (*5, 21, 36*). We generated B16F10 melanoma to constitutively express mouse VEGF-C (B16/VEGF-C) and VEGF-C expression was confirmed in vitro by qPCR, ELISA, Western Blot, and by immunostaining (Fig. 1a-c, Suppl. Fig. 1a-b). B16/VEGF-C cells secreted 33 kDa VEGF-C protein, which binds VEGFR-3 but not VEGFR-2 (*29, 30*) (Fig. 1c). B16/EV cells, transduced with an empty vector (B16/EV) showed no detectable expression of VEGF-C mRNA or protein (Fig. 1a-c, Suppl. Fig. 1a-b).

**Fig 1.**
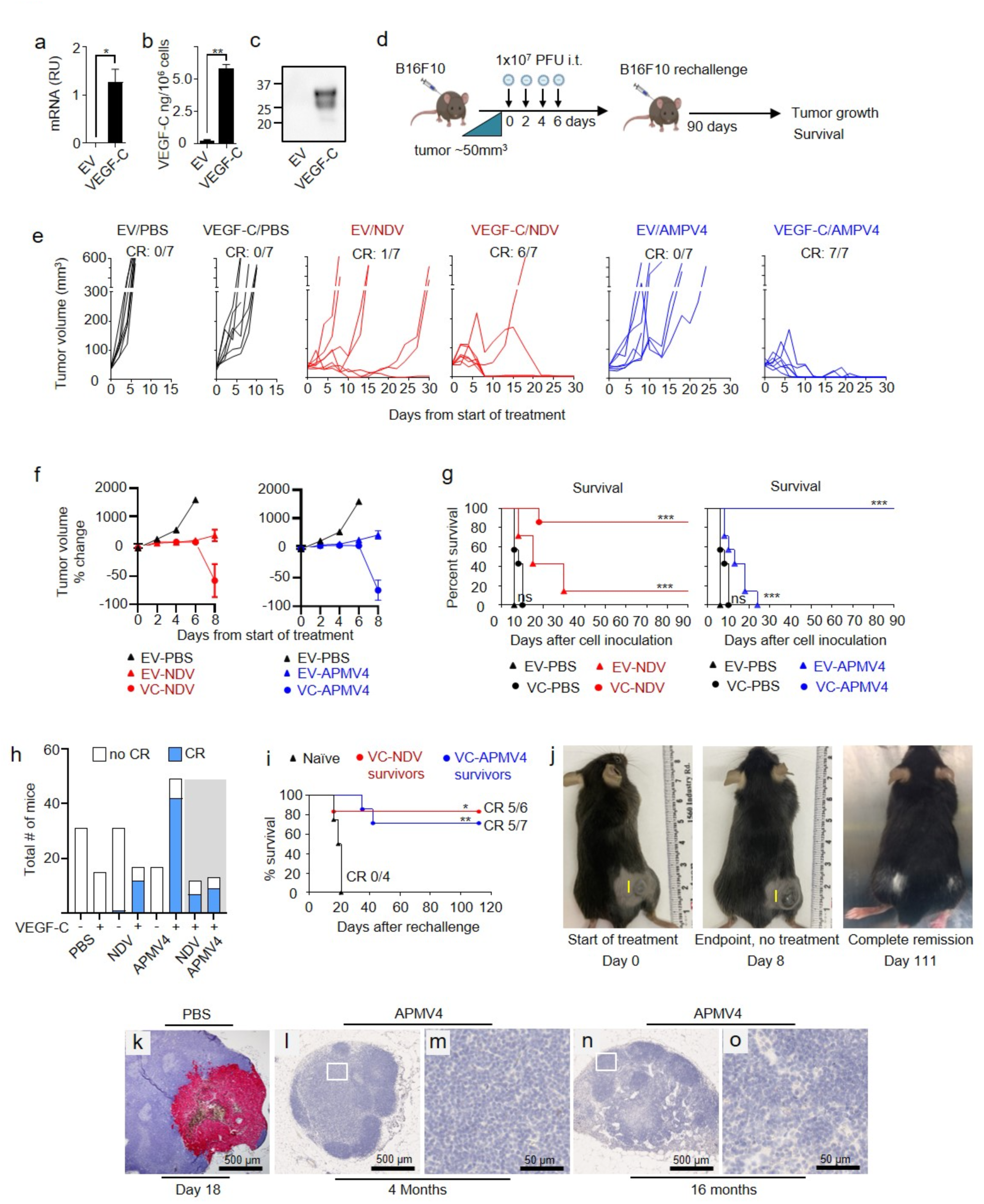
VEGF-C enhances the anti-tumor efficacy of APMVs in the B16F10 tumor model. (a-c) VEGF-C expression by B16F10 melanoma cells, transduced with mVEGF-C or an empty vector (EV), determined by qPCR (a), ELISA (b), and Western blot (c). Error bars represent mean ± SEM, n=2. Statistics: unpaired t-test *p*=0.0417 (a) and *p*=0.029 (b). (d) Experimental design: Mice were injected with 3×10^5^ B16F10/EV or B16F10/VEGF-C cells. Treatment began when tumors reached ∼50mm^3^. NDV, APMV4 or PBS were administered intratumorally four times, every two days, and mice were monitored for tumor growth and survival. (e) B16/EV and B16/VEGF-C tumor growth upon treatments as indicated. Individual tumor growth curves are shown. (f) Tumor volume percentage change relative to the start of treatment over the first eight days. Error bars represent mean ± SEM, n=7 (g) Survival of mice bearing B16/EV or B16/VEGF-C tumors, treated with PBS, NDV, or APMV-4, as indicated. Statistical analysis: Log-rank (Mantel-Cox) test, n=7. (h) Cumulative data from all experiments showing total number of mice per experimental condition, number of mice in each group achieving complete remission (blue), and those not achieving complete remission (white). Gray shaded area indicates rechallenge experiments. (i) Survival of mice rechallenged with B16F10/EV cells 90 days after achieving complete remission (CR) following treatment of B16/VEGF-C tumors with NDV or APMV4. Statistical analysis: Log-rank (Mantel-Cox) test. (j) Representative images of mice at three time points: left, start of the treatment; center, humane end-point untreated mouse; right, mouse in complete remission after NDV treatment of B16F10/VEGF-C tumors and subsequent rechallenge. White patch of hair on the site of the first regressed tumor (right hind limb) and on the site of rechallenge (left hind limb). (k-o) Melan-A staining for detection of B16 melanoma metastases. Representative images of sentinel (inguinal) LNs of mice bearing B16F10/VEGF-C tumors, treated with PBS (n=18, 2 experiments) or APMV4 (n=23, 4 experiments) at indicated time-points after the start of treatment. In vivo experiments were independently conducted 2-8 times. \**p*<0.05 \*\**p*<0.01 \*\*\**p*<0.001, ns = not significant. Scale bars: i, 5 mm; k, m, o, 500 μm; l, n, 50 μm.

B16/VEGF-C and B16/EV tumors (∼50mm^3^) were treated with NDV or APMV-4 by intratumoral injection every two days, for four doses total (1×10^7^ PFU each) (Fig. 1d). NDV delayed B16/EV tumor growth (5/7), with 14% mice in complete remission (1/7), while triggering complete remission in 87% of mice with B16/VEGF-C tumors (6/7) (Fig. 1e). APMV-4 treatment of B16/EV tumors also significantly delayed tumor growth (5/7), but did not induce CR. However, when tumors expressed VEGF-C, APMV-4 treatment led to complete remission in 100% of the mice (7/7) (Fig. 1e). We observed a transient increase in tumor volume after two treatments, prior to tumor regression (Fig. 1e). This is consistent with the previous studies of immune checkpoint inhibition and has been correlated with immune cell infiltration and inflammation (*15*). Tumors regressed rapidly; by day eight following treatment with APMV-4 or NDV, 71% of tumors had fully regressed (Fig. 1f). Mice were monitored post-treatment with either NDV or APMV-4 for up to 2 years and 4 months (843 days), during which no tumor recurrence was observed, indicating complete tumor eradication (Fig. 1g, Table 1). Cumulative data from eight independent experiments show that APMV-4 is highly effective on tumors expressing VEGF-C, leading to complete remission (CR) in 86% of mice (n=49), compared to 0% CR when VEGF-C was not present (n=17) (Fig. 1h, Table 1). APMV-4 was more efficient in eliminating B16/VEGF-C tumors than NDV (86% vs. 71%, respectively) (Fig. 1h, Table 1).

**Table 1.** Complete response rates upon treatment of B16/EV or B16/VEGF-C tumors with NDV or APMV4. Number of tumor-free surviving mice over total number of mice in an experiment is indicated. Data are shown from ten independent experiments. CR, complete response.

|  | PBS |  | NDV |  | rechallenge | APMV4 |  | rechallenge |
| --- | --- | --- | --- | --- | --- | --- | --- | --- |
| Exp # | EV | VEGF-C | EV | VEGF-C | EV | EV | VEGF-C | EV |
| 1 | 0/9 | 0/8 | 0/10 | 6/10 | 2/6 | 0/10 | 7/9 | 4/6 |
| 2 | 0/7 | 0/7 | 1/7 | 6/7 | 5/6 | 0/7 | 7/7 | 5/7 |
| 3 | 0/8 |  | 0/10 |  |  |  |  |  |
| 4 | 0/7 |  | 0/7 |  |  |  |  |  |
| 5 |  |  |  |  |  |  | 9/9 |  |
| 6 |  |  |  |  |  |  | 2/5 |  |
| 7 |  |  |  |  |  |  | 4/4 |  |
| 8 |  |  |  |  |  |  | 5/5 |  |
| 9 |  |  |  |  |  |  | 3/5 |  |
| 10 |  |  |  |  |  |  | 5/5 |  |
| CR | 0/31 | 0/15 | 1/34 | 12/17 | 7/12 | 0/17 | 42/49 | 9/13 |
| %CR | 0% | 0% | 3% | 71% | 58% | 0% | 86% | 69% |

### Surviving mice are metastasis free and protected from tumor development upon re-challenge

To assess the durability of response to NDV or APMV-4 therapy, mice in complete remission (CR) were re-challenged with B16/EV tumor cells 90 days post-treatment, by injecting tumor cells into the contralateral flank. As expected, naïve control mice rapidly developed tumors, reaching maximum size within 20 days (Fig. 1i). In contrast, most mice in CR were protected from developing tumors after re-challenge and remained tumor-free during the observation period of up to 546 days (∼18 months) (Fig. 1i, j, Table 1). Mice frequently displayed vitiligo at tumor rejection sites, consistent with local immune activation (Fig. 1j), as reported previously for checkpoint blockade (*37*). Cumulative protection rates post re-challenge were 69% (n=13) in APMV-4 and 58% (n=12) in NDV-treated group (Fig. 1h, Table 1). These findings demonstrate that NDV and APMV-4 treatments consistently induce robust and sustained anti-tumor immunity when VEGF-C is present in tumors.

To assess effects of APMV-4 treatment on metastases, we immunostained lymph nodes from B16/VEGF-C tumor-bearing mice for MelanA (Fig. 1k-o, Table 2). By day 21 after tumor inoculation, all mice in control group developed macrometastases in sentinel LNs (Fig. 1k). Strikingly, in sentinel LNs of APMV-4-treated mice metastases were not detectable for the entire observation period, ranging from 4-18 months (Fig. 1k-o, Table 2). These results demonstrate that APMV-4 treatment of B16 tumors expressing VEGF-C eliminates lymph node metastases, indicating that metastasis-promoting effects of VEGF-C can be reversed in the context of immune activation, such as during viral therapy.

**Table 2.** Cumulative incidence of lymph node metastasis. Number of mice with sentinel LNs positive for metastasis. Each mouse had one tumor and one sentinel lymph node.

|  | PBS | APMV4 |  |
| --- | --- | --- | --- |
| Exp # | VEGF-C |  | Days after treatment |
| 1 | 10/10 |  |  |
| 2 | 8/8 |  |  |
| 3 |  | 0/4 | 120 |
| 4 |  | 0/5 | 454 |
| 5 |  | 0/5 | 843 |
| 6 |  | 0/9 | 751 |
| # LN w mets | 18/18 | 0/23 |  |
| % LN w mets | 100% | 0% |  |

### VEGF-C-expressing tumors are enriched in T cells

To assess the effects of VEGF-C on the tumor microenvironment, we carried out immunostaining for lymphatic and blood vasculature, as well as for the key immune cell subsets (Fig. 2a). Tumors were collected 12h after the second treatment, when tumor sizes were comparable across all treatment groups, and just prior to the onset of regression (Suppl. Fig. 2a).

**Fig 2.**
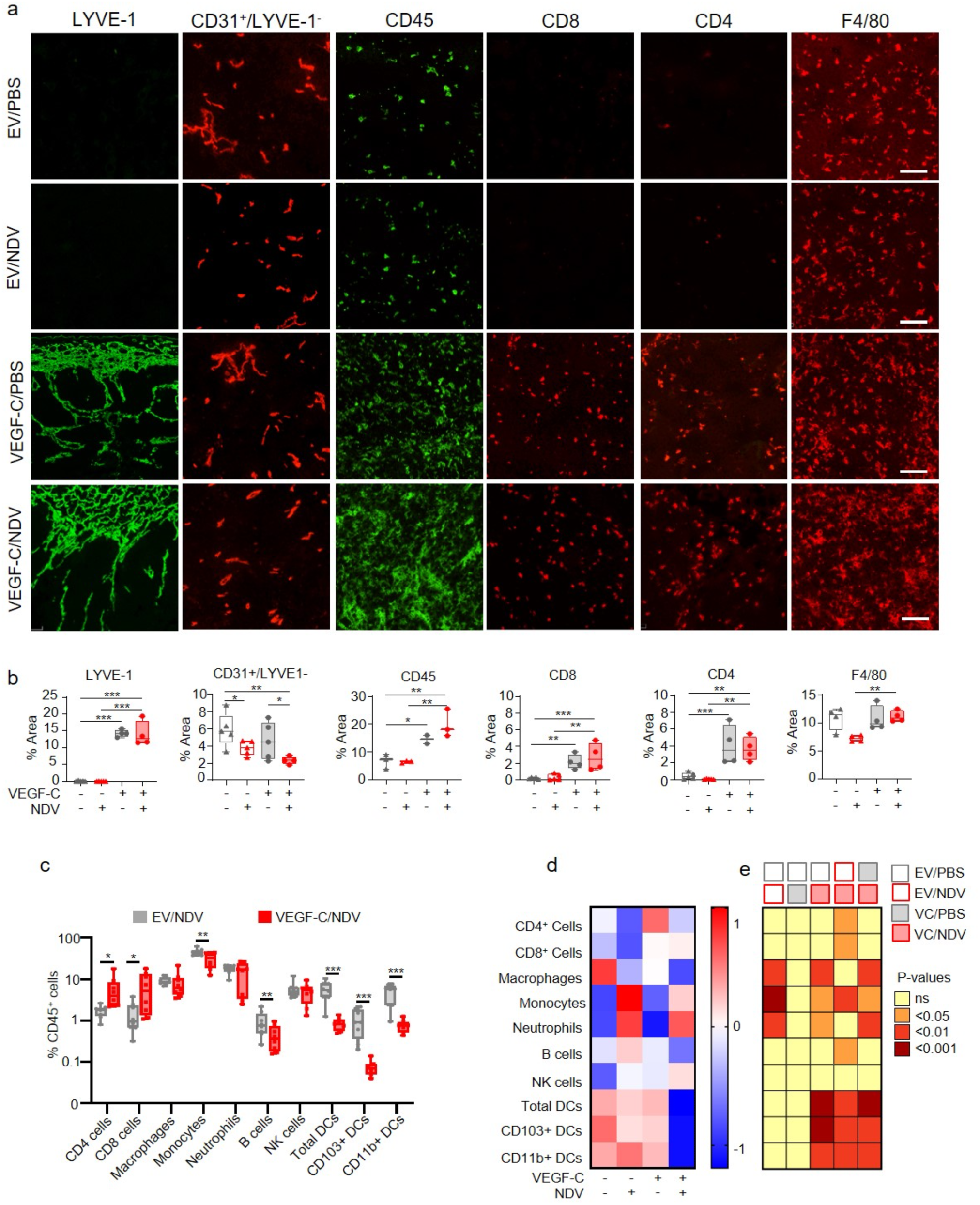
Effects of VEGF-C and NDV on tumor vasculature and immune cells. (a, b) Immunofluorescent staining (IF) of B16/EV and B16/VEGF-C tumors treated with PBS or NDV, for lymphatics (LYVE-1), blood vessels (CD31+/LYVE-1-), and immune cells, as indicated. Representative images (a), and quantification of immunostaining (b). Box-and-whiskers plots show individual values with mean ± SEM, and whiskers spanning the full range; EV/PBS, n=5; EV/NDV, n=5; VEGF-C/PBS, n=4; VEGF-C/NDV, n=4. (c) Immunophenotyping of NDV-treated B16/EV and B16/VEGF-C tumors by spectral flow cytometry. Frequencies of distinct myeloid and T cell subtypes as a percentage of total CD45 cells are shown. Box-and-whiskers plots show individual values with mean ± SEM, and whiskers spanning the full range; EV/NDV, n=8; VEGF-C/NDV, n=8. Outliers were removed based on ROUT analysis. Q=0.5%. (d) Heatmap displaying z-score values of various immune cell populations in B16 tumors, as indicated. (e) Heatmap displaying p-values for the multiple comparisons between the populations in (d). Statistical analysis with one-way ANOVA multiple comparisons Fisher LSD test. *p < 0.05, **p < 0.01, ***p < 0.001, ns = not significant. Scale bars: 100 µm.

B16/EV tumors did not express VEGF-C (Fig. 1a-c) and showed no intratumoral or peritumoral lymphangiogenesis, as indicated by LYVE-1 staining (Fig. 2a, b). NDV-treated B16/EV tumors similarly lacked lymphatics in or around the tumors (Fig. 2a, b). In contrast, VEGF-C expression induced tumor lymphangiogenesis, as expected (Fig. 2a, b), with higher lymphatic densities at the tumor periphery than in the center, and comparable distribution in PBS- and NDV-treated B16/VEGF-C tumors (Fig. 2a, b). VEGF-C also promoted lymphangiogenesis and dilation of lymphatics in overlying skin, while in the skin adjacent to B16/EV tumors, lymphatics appeared normal in density and morphology (Suppl. Fig. 2b). NDV treatment alone did not induce lymphangiogenesis or alter the appearance of lymphatics in tumors or skin within 2.5 days from the onset of treatment (Fig. 2a, b; Suppl. Fig. 2). Angiogenesis was not induced by VEGF-C. However, NDV decreased blood vessel area in tumors by decreasing both, their size and density (Fig. 2a, b).

VEGF-C markedly increased CD45^+^ cells in tumors, whereas NDV did not within 2.5 days from the onset of treatment (Fig. 2a, b; Suppl. Fig. 2). Moreover, VEGF-C expression was associated with high densities of CD8^+^ and CD4^+^ T-cells in tumors, while VEGF-C-negative tumors were nearly devoid of T cells at this time-point (Fig. 2a, b; Suppl. Fig. 2c). Densities of F4/80^+^ macrophages were high across all groups, with no changes due to VEGF-C or NDV (Fig. 2a, b). Conversely, Ly6G^+^ neutrophils were present peritumorally, but absent within tumors across all treatment groups (Suppl. Fig. 2c).

VEGFR-3, the key VEGF-C receptor, was detected on all LYVE-1^+^ lymphatic vessels in B16/VEGF-C tumors by immunostaining (Suppl. Fig. 3), with no VEGFR-3 detected on other cell types, and no changes in the levels or pattern of VEGFR-3 expression upon NDV treatment (Suppl. Fig. 3). Taken together, these findings demonstrate a positive correlation between lymphangiogenesis and T-cell densities and suggest that VEGF-C promotes T cell expansion indirectly, by activating VEGFR-3 signaling on lymphatic endothelial cells (LECs).

### Immune profiling of tumors reveals unique changes of immune cell composition associated with complete response

To investigate mechanisms underlying tumor growth delay (EV/NDV) or complete regression (VEGF-C/NDV), we profiled immune cell phenotypes and activation states in the tumor microenvironment. High-dimensional spectral flow cytometry was performed on tumors collected 12h after the second NDV treatment (Suppl. Fig. 2a), using markers for myeloid and lymphoid lineages (Table 3, Panel 1). Given the high complete response (CR) rate in B16VEGF-C/NDV-treated animals (71%), changes in immune cell subsets could be linked to tumor regression with high confidence.

**Table 3.** Markers used in spectral flow cytometry.

| **Panel 1: Tumor**<br>Myeloid and lymphoid<br>Cell surface | **Panel 2: Tumor**<br>Lymphoid<br>Cell surface | **Panel 3: Tumor**<br>Lymphoid<br>Intracellular | **Panel 4: Lymph Nodes**<br>Myeloid and lymphoid<br>Cell surface |
| --- | --- | --- | --- |
| B220 | CD62L | B220 | B220 |
| CCR2 | CD25 | CD3 | CD103 |
| CD103 | CD28 | CD4 | CD11b |
| CD11b | CD3 | CD45 | CD11c |
| CD11c | CD4 | CD49b | CD3 |
| CD45 | CD44 | CD8 | CD44 |
| CD49b | CD45 | Foxp3 | CD45 |
| CD64 | CD49b | GrzmB | CD49b |
| CD8 | CD8 | IFNγ | CD62L |
| CD80 | CTLA-4 | Live Dead | CD64 |
| CD83 | CXCR3 | NK1.1 | CD8 |
| CD86 | KLRG1 | TNFα | CD83 |
| F4/80 | Live Dead |  | CD86 |
| Live Dead | PD-1 |  | Foxp3 |
| Ly6C | TIM-3 |  | Live Dead |
| Ly6G |  |  | Ly6C |
| MHCII |  |  | Ly6G |
| PD-L1 |  |  | MHCII |
|  |  |  | PD-L1 |

Analysis of B16/VEGF-C NDV-treated tumors by manual gating showed an increase in relative frequencies of CD4^+^ and CD8^+^ T cells, and a decrease in dendritic cells (DCs), monocytes and B cells associated with CR (Fig. 2c-e; Suppl. Fig. 4). CD103^+^ DCs, which are a key antigen-presenting cells (APCs) priming tumor-specific CD8+ T cells(*38*), as well as CD11b^+^ DCs, showed reduced frequencies only in NDV-treated B16/VEGF-C tumors (Fig. 2c-e, Suppl. Fig. 5a). These findings suggest enhanced DC mobilization to LNs following NDV treatment of VEGF-C-expressing tumors, which is consistent with the previous study demonstrating that mobilizing DCs improved NDV-based virotherapy against B cell lymphoma (*39*).

NDV treatment of B16/EV tumors, which induced partial tumor regression, increased monocyte and neutrophil frequences compared to PBS controls (Fig. 2d, e; Suppl. Fig. 5a). Neutrophils were localized almost exclusively at the tumor periphery, as demonstrated by immunostaining (Supp. Fig. 5b). NDV treatment of both B16/EV and B16/VEGF-C tumors showed a trend toward increased NK cell frequencies compared to PBS controls, but it was not statistically significant, likely due to the low NK cell abundance relative to other immune cell subtypes (Fig. 2d, e; Suppl. Fig. 5a). These data highlight several key differences in immune cell composition in response to NDV treatment, depending on the presence or absence of VEGF-C.

### VEGF-C and NDV cooperate to shift the tumor immune landscape toward effector CD8⁺ T cells

Given the increase in CD8^+^ and CD4^+^ T cells in VEGF-C-expressing tumors, we analyzed T cell phenotypes associated with partial and complete anti-tumor responses. Analysis of T-cell surface markers (Table 3, Panel 2) by manual gating revealed a significant increase in CD25⁺PD-1⁺CD8⁺ T cells in VEGF-C/NDV tumors, correlating with complete response (Fig. 3a, b). VEGF-C alone also increased this subset, while NDV showed a trend (Fig. 3b). In contrast, CD44⁺PD-1⁺CD8⁺ T cell frequences were not significantly changed across conditions (Fig. 3c, Suppl. Fig. 6a), highlighting CD25 as a distinguishing marker associated with CR. CD25^+^CD8^+^ T cells were mainly expanded by NDV, and this was further enhanced by VEGF-C (Suppl. Fig. 6b-e). VEGF-C, however, strongly increased PD-1 expression on this subset (Suppl. Fig. 6f, g). All CD8^+^CD25^+^ T cells expressed CD44^+^, suggesting that they are Ag-experienced (Suppl. Fig. 6h). There was in increase in CTLA-4^+^ CD8⁺CD25⁺PD-1⁺ T cells upon NDV treatment, but they lacked TIM-3 (Fig. 3d). Notably, frequencies of CD4^+^Foxp3^+^ Tregs were significantly decreased and the ratio of CD25^+^PD-1^+^CD8^+^ T cells to Tregs was increased only in VEGF-C/NDV tumors (Fig. 3e, f). Taken together, these data demonstrate robust expansion of CD25^+^PD-1^+^ effector CD8^+^ T cells and reduction of Tregs selectively in NDV-treated B16/VEGF-C tumors, suggesting a key role of these subsets in tumor eradication.

**Fig 3.**
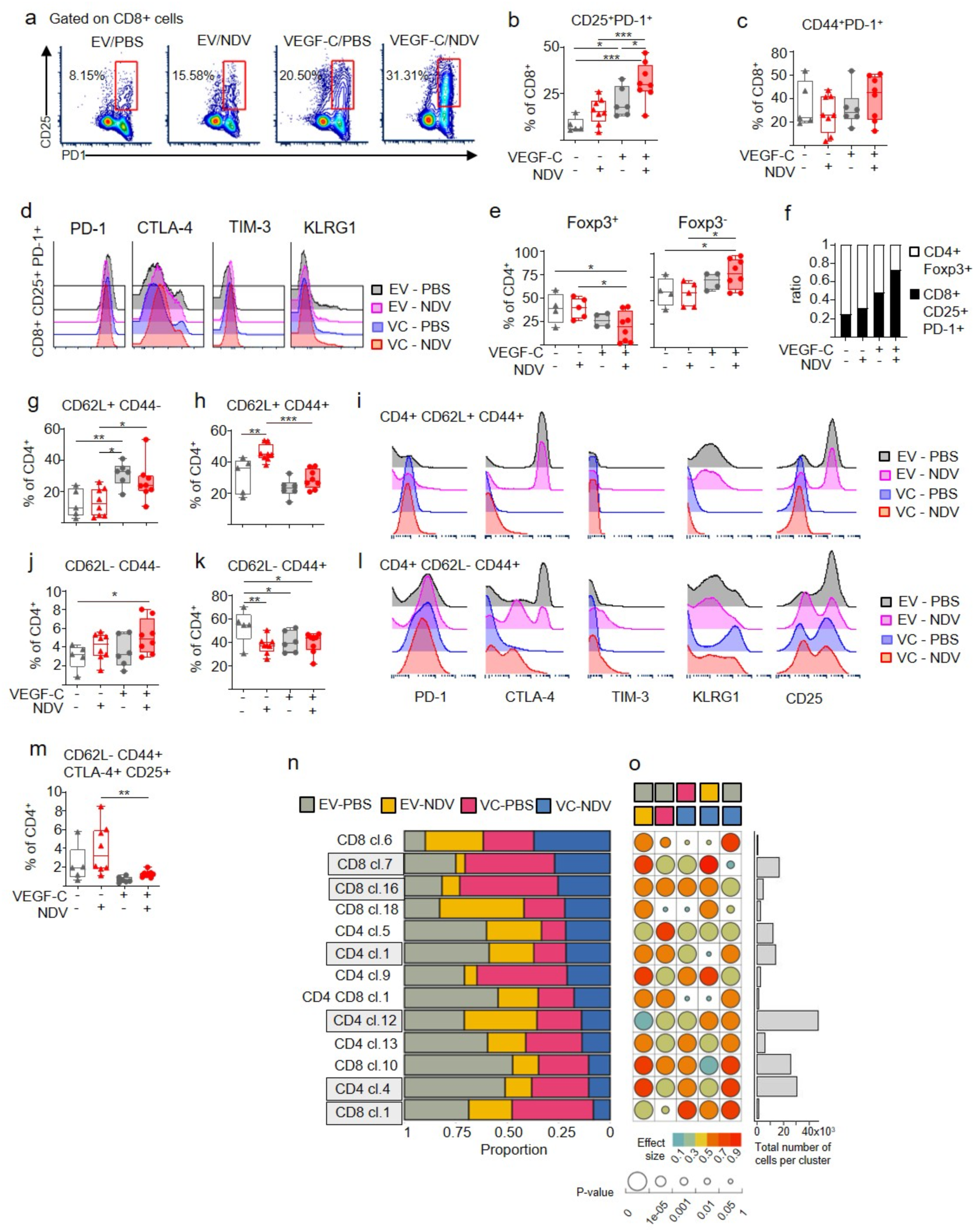
Effects of VEGF-C and NDV on T cell phenotypes. (a-p) Spectral flow cytometry analysis of T-cell subsets in B16/EV and B16/VEGF-C tumors treated as indicated. (a-c) Frequencies of CD25^+^PD-1^+^ CD8^+^ T cells; contour plots (a) and Box-and-whiskers plots (b). (c) Frequences of CD44^+^PD-1^+^ CD8^+^ T cells. (d) Histograms of checkpoint receptors on CD25^+^PD-1^+^ CD8^+^ T cells across treatments. (e) Frequencies of Foxp3^+^ and Foxp3^-^ CD4^+^T cells. (f) Ratios of CD25^+^PD-1^+^ CD8 effector T cells to Foxp3^+^ Tregs across treatments as indicated. (g-m) Frequencies of various CD4^+^ T cell subsets (g,h,j,k, m) and histograms for immune check points (i, l) as indicated. Box-and-whiskers plots show individual values with mean ± SEM, and whiskers spanning the full range. EV/PBS, n=5; EV/NDV, n=8; VEGF-C/PBS, n=6; VEGF-C/NDV, n = 8. Statistical analysis with one-way ANOVA multiple comparisons Fisher LSD test. *p < 0.05, **p < 0.01, ***p < 0.001, not indicated = not significant. (n) Stacked barplots showing T cell cluster proportions in tumors across treatments. Different experimental groups are represented by their respective colors. Only T cell clusters which showed significant difference between any of the conditions are shown. Most significantly changed T cell clusters are shaded in gray. (o) Heatmap showing p-values (dot size) and effect size (Cohen’s W; dot color) of the Chi-square goodness-of-fit test for pairwise comparison of cell cluster proportions across treatments. Only clusters with over 100 cells, medium or large overall effect size and a Chi-square p-value of less than 0.05 in comparisons of different treatments are depicted. The bar plot on the far right displays the total number of cells in each cluster.

VEGF-C exhibited potent effects on CD4^+^ T cells. VEGF-C increased frequencies of naïve CD4^+^ T cells (CD62L^+^CD44^-^) (Fig. 3g), but it reduced CD62L^+^CD44^+^ CD4^+^ T cells, which were abundant in B16F10 tumors, and expanded with NDV (Fig. 3h). In untreated tumors, CD62L^+^CD44^+^ CD4^+^ T cells were positive for CD25, CTLA-4, and KLRG1, and this population was increased in frequencies with NDV (Fig. 3h, i). However, in VEGF-C expressing tumors, CD25, CTLA-4, and KLRG-1 were significantly reduced on CD62L^+^CD44^+^ CD4^+^ T cells (Fig. 3i), suggesting reduced sensitivity to IL-2 and reduced inhibitory signaling. VEGF-C also markedly decreased CD62L^-^CD44^+^CTLA-4^+^ CD4^+^ T cells (Fig. 3l). It also increased KLRG1^+^ subset, but did not change PD-1 across conditions (Fig. 3l). Either NDV or VEGF-C alone reduced frequences of CD25^+^CD62L^-^CD44^+^ CD4^+^ T cells (Fig. 3l), whereas VEGF-C potently decreased CTLA-4^+^CD25^+^ subset (Fig. 3m). Decrease in CTLA-4^+^CD25^+^ CD4^+^ T cell subsets, likely Tregs, suggests reduced capacity of these cells to suppress immune responses.

Unsupervised bioinformatic analysis separately identified three CD8^+^ T cell clusters with significantly changed frequences in VEGF-C/NDV group compared to EV/NDV. Specifically, effector CD8^+^ T cell clusters 7 and 16, both CD28^+^CD44^+^PD-1^+^, were the most enriched populations, whereas CD8^+^ T cell cluster 1 (CD28^+^CD44^+^ CTLA-4^+^TIM-3^+^), likely early exhausted cells, were decreased (Fig. 3 n, o; Suppl. Fig. 7). Effector CD8^+^ T cell cluster 7 was also the most abundant of all CD8 clusters affected by treatment (Fig. 3n, o). The number of CD4⁺ T cells affected by treatments was approximately double that of CD8⁺ T cells (CD4:CD8 ratio = 2.18:1). CD4⁺ T cell clusters decreased with NDV/VEGF-C were all CTLA-4⁺; (Cluster 4 and 12, CD44⁺CD28⁺CTLA-4⁺) and Cluster 1 (CD44⁺CD28⁺CTLA-4⁺PD-1^+^TIM-3^+^). Cluster 1 was most abundant CD4⁺ T cell subset among those affected by treatment. Taken together, VEGF-C and NDV combined drive a strong anti-tumor immune response by reducing immunosuppressive or exhausted cells and expanding effector T cells. Since VEGF-C alone increases total CD8^+^ T cells, this results in increased total T cell effectors in VEGF-C/NDV tumors, correlating with tumor eradication.

### T cells and NK cells are highly activated in VEGF-C-expressing tumors treated with NDV

We next analyzed the activation status of NK, CD8^+^, and conventional CD4^+^ T cells, by assessing the expression of IFNγ, TNFα and granzyme B (GzmB) using spectral flow cytometry (Table 3, Panel 3). Manual gating revealed strong activation of NK and CD8^+^ T cells, with NK cells showing very high activation levels (Fig. 4a. b, Suppl. Fig. 8b-d). NDV markedly increased IFNγ^+^ NK cells in tumors and this was further enhanced by VEGF-C (Fig. 4b). NDV also triggered a 10-fold increase in GzmB^+^ NK cells independent of VEGF-C, indicating enhanced cytotoxic capacity of NK cells. Conversely, VEGF-C, but not NDV, increased TNFα^+^ NK cells (Fig. 4b), most of which co-expressed IFNγ^+^ (Suppl. Fig. 7d).

**Fig 4.**
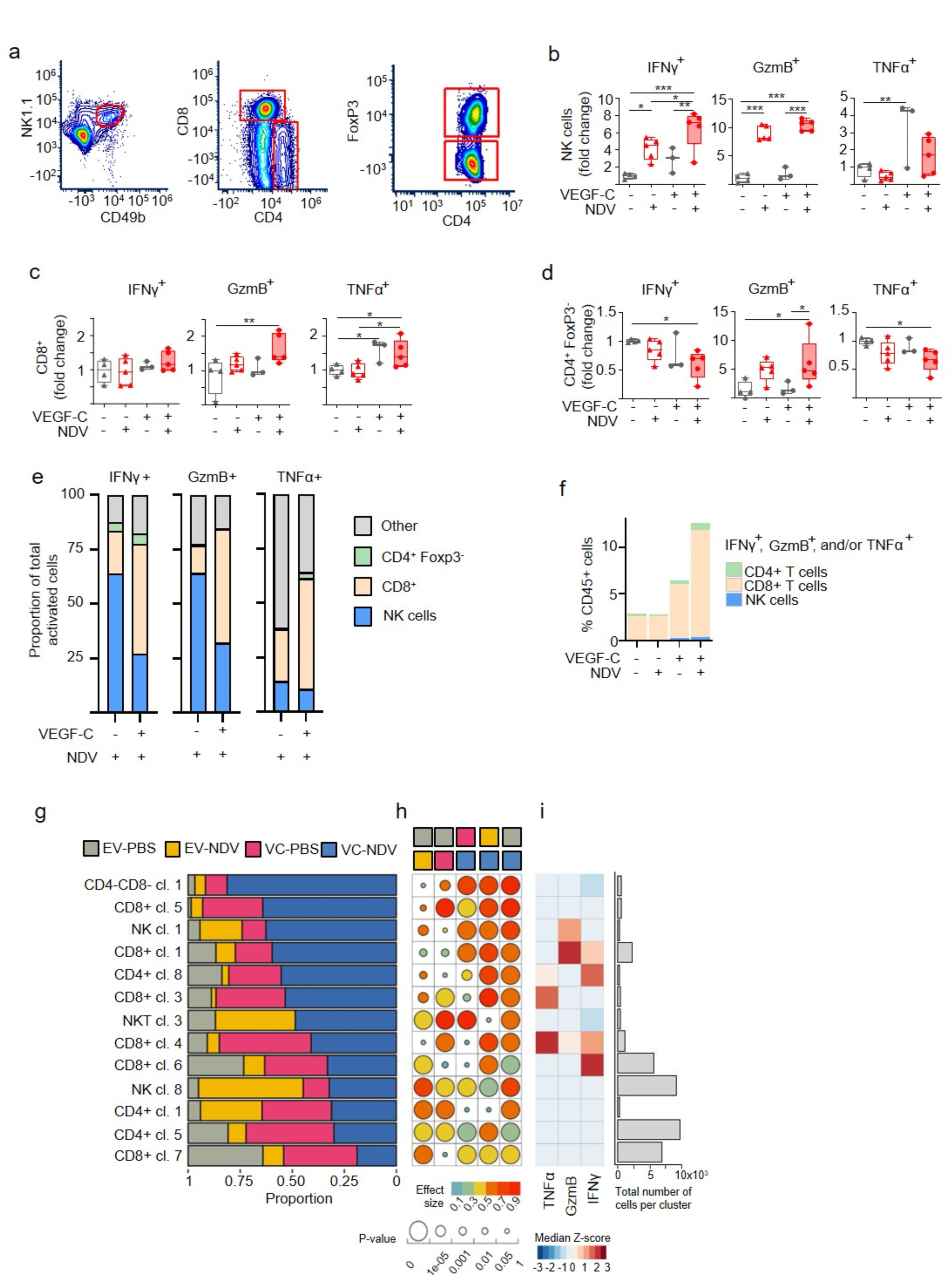
Effects of VEGF-C and NDV on T cell and NK cell activation in B16F10 tumors. (a-i) Spectral flow cytometry analysis of T cell and NK cell activation. (a) Gating strategy for NK cells, CD8^+^, CD4^+^, and CD4^+^ Foxp3^+^ T cells. (b-d) Increase or decrease (fold change from EV/PBS control group) of NK cells (b), CD8^+^ T cells (c), and CD4^+^ conventional T cells (d), positive for IFNγ, granzyme B (GzmB), or TNFα upon treatments as indicated. Box-and-whiskers plots show individual values with mean ± SEM, and whiskers spanning the full range. EV/PBS, n=4; EV/NDV, n=5; VEGF-C/PBS, n=3; VEGF-C/NDV, n=5. Statistical analysis with one-way ANOVA multiple comparisons Fisher LSD test. (e) Proportion of cells positive for IFNγ, GzmB or TNFα in NDV-treated B16/EV and B16/VEGF-C tumors. (f) Cumulative percentages of activated CD4^+^ T cells, CD8^+^ T cells, and NK cells as indicated. (g) Stacked barplots showing cell cluster proportions in tumors across treatments. Different experimental groups are represented by their respective colors. Only T cell clusters which showed significant difference between any of the conditions are shown. (g) Heatmap showing p-values (dot size) and effect size (Cohen’s W; dot color) of the Chi-square goodness-of-fit test for pairwise comparisons of cell cluster proportions across treatments. Only clusters with over 100 cells, medium or large overall effect size and a Chi-square p-value of less than 0.05 in comparisons of different treatments are depicted. (i) Heatmap showing the median of z-score normalized mean marker expression values for cell clusters identified based on expression of TNF-α, GrB and/or IFN-γ. The bar plot on the far right displays the total number of cells in each cluster. *p < 0.05, **p < 0.01, ***p < 0.001, not indicated = not significant.

CD8^+^GrB^+^ T cells, which are potent cytotoxic T cells, were significantly increased only in VEGF-C/NDV tumors (Fig. 4c). VEGF-C increased TNFα^+^ CD8^+^ T cells which were also IFNγ^+^ (Fig. 4c, Suppl. Fig. 8d), while NDV alone had no effect on CD8^+^ T cell activation at the time-point examined (Fig. 4c). Conventional CD4^+^ T cells in VEGF-C/NDV tumors showed an NDV-driven increase in GzmB, and a decrease in IFNγ^+^ and TNFα^+^ CD4^+^ T cells (Fig. 4d, Suppl. Fig. 8d). Taken together, these data demonstrate enhanced activation of NK, CD8^+^ and CD4^+^ T cells in NDV-treated VEGF-C-expressing tumors, correlating with complete response.

Analysis of the proportional abundance of activated CD45^+^ cells demonstrated that NDV treatment mainly activated NK cells, as indicated by IFNγ and GzmB expression. In the presence of VEGF-C, however, the ratio of activated CD8^+^ T cells to NK cells increased in NDV-treated tumors (Fig. 4e, Suppl. Fig. 8e, f). VEGF-C alone had minimal or no impact on proportions of activated cells in untreated tumors (Suppl. Fig. 8e, f), consistent with the observation that VEGF-C had no anti-tumor effects in the absence of immune activation with the virus. Finally, relative proportion of activated effector cells among all CD45+ cells increased ∼5-fold in VEGF-C/NDV tumors and ∼2-fold in untreated B16/VEGF-C tumors (Fig. 4f). Together, these data demonstrate a marked accumulation of activated effector cells, particularly CD8^+^ T cells, in VEGF-C/NDV tumors.

Unsupervised bioinformatic analysis separately identified six clusters of T cell and NK cell populations with distinct expression patterns of effector molecules, including three clusters of polyfunctional T cells (Fig. 4g-i, Suppl. Fig. 8g). Four CD8^+^ T cell, one CD4^+^ T cell and one NK cell cluster displayed enhanced production of GgzmB, IFNγ and/or TNFα. Notably, all six clusters were significantly expanded in VEGF-C/NDV group compared to EV/NDV (Fig. 4g-i). In summary, these data show that NDV combined with VEGF-C strongly activates CD8^+^ T cells, conventional CD4^+^ T cells and NK cells, and that the expansion of activated effector cells is linked to tumor eradication.

### T cells establish close contact with tumor lymphatic endothelium

We observed expansion of T-cell compartment in VEGF-C expressing tumors, with maximal activation following NDV treatment in the presence of VEGF-C. To better understand the spatial relationship between lymphatics and T cells, we performed immunostaining on B16/VEGF-C tumor samples collected 12h following the second dose of NDV treatment (Fig. 5). High lymphatic vessel densities in tumors correlated with high CD8^+^ and CD4^+^ T cell densities (Fig 5a-j). Analysis of lymphatic vessel distribution demonstrated that lymphatic densities were high at the tumor periphery (Fig. 5a, f) and sparse in the tumor center (Fig. 5c, h). Accordingly, T cell densities were high at the tumor periphery, and sparse in the tumor center, mirroring lymphatic vessel distribution (Fig. 5a-j). Most T cells localized near lymphatics, suggesting direct interaction with tumor LECs (Fig. 5k), and z-stack confocal microscopy and fluorescence intensity analysis confirmed direct contact between CD8^+^ T cells and LECs (Fig. 5l, m).

**Fig 5.**
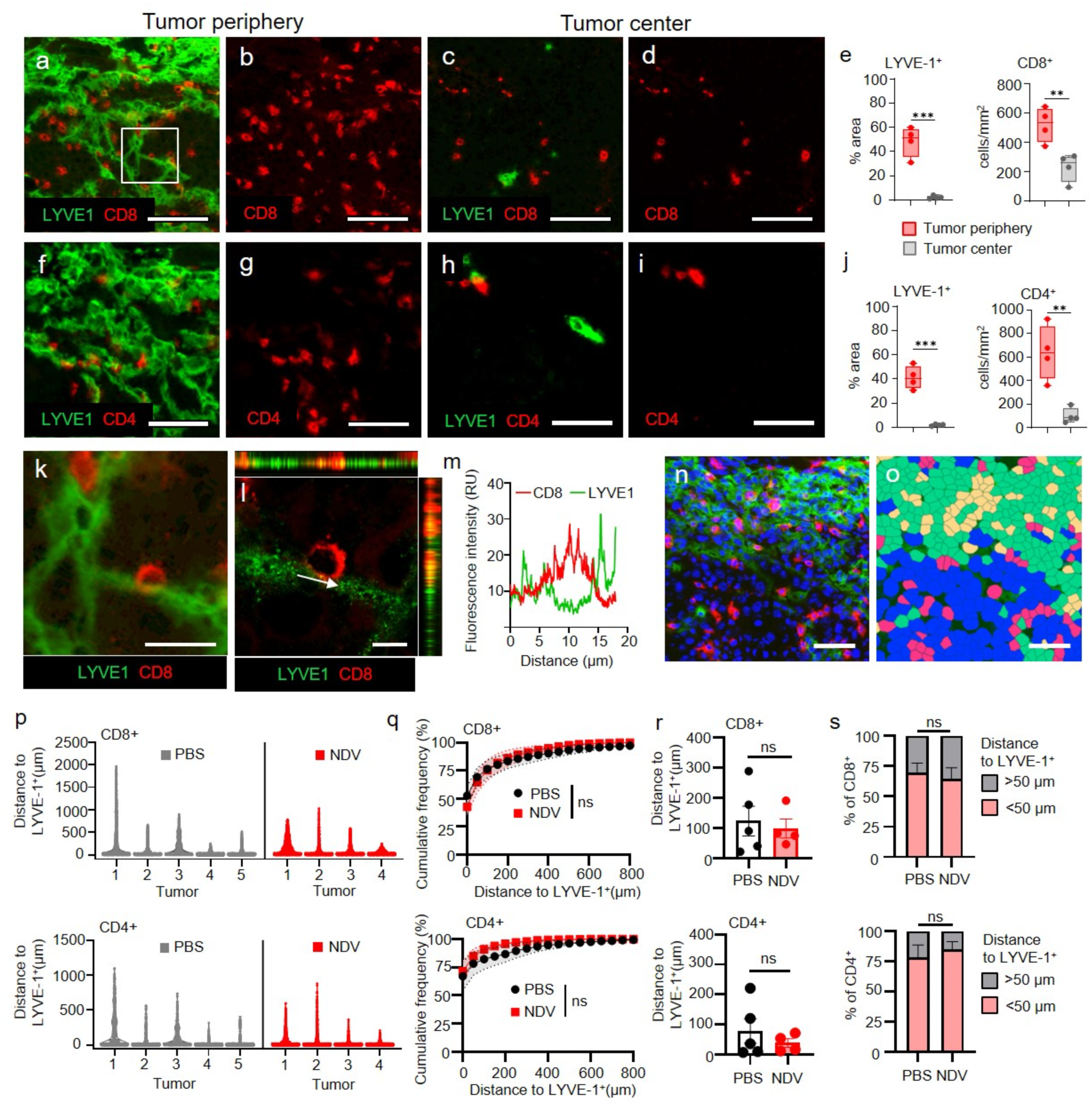
T-cells localize near lymphatic vessels in tumors. (a-d, f-i) NDV-treated B16/VEGF-C tumors immunostained for LYVE-1 (lymphatics, green), and CD8 or CD4, as indicated. Tumor areas with dense and sparse lymphatics are shown. (e, j) Quantification of lymphatic vessel densities and T cells in tumor areas with high and low lymphatic densities, n=4. Box-and-whiskers plots show individual values with mean ± SEM, and whiskers spanning the full range. (k, l) High magnification image (k) and a z-stack (l) of a boxed area in 5a, showing lymphatic endothelium and CD8^+^ T cell in close contact. (m) Quantification of the fluorescence intensity in the indicated area, in the direction of the arrow (l). (n, o) Immunofluorescent staining of a representative B16/VEGF-C NDV-treated tumor for CD8^+^ (red), LYVE-1^+^ (green) and DAPI (blue) (n), and a corresponding Qu-Path overlaying mask used for proximity analyses (o). (p-s) Proximity analyses of CD8^+^ and CD4^+^ T cells to tumor lymphatics in B16/VEGF-C tumors treated with PBS or NDV. Quantification of the distances from each T cell to the nearest LYVE-1^+^ lymphatic vessel (p); Cumulative frequencies of T cells within various distances to the nearest LYVE-1^+^ lymphatic vessel. Shaded area indicates STD (q); Average distance of T cells to the nearest LYVE-1^+^ vessel (r); T cell frequences within < 50µm or > 50µm of the nearest LYVE-1^+^ vessel. Statistical analysis: two-tailed student t-test. **p < 0.01, ***p < 0.001, ns = not significant. Scale bars: a-d, f-i: 100 µm; k: 25 µm, l: 10 µm; n, o: 50 µm.

Proximity analysis revealed that the majority of CD8^+^ T cells in tumors were in close proximity to lymphatics, irrespective of NDV treatment (Fig. 5n-s). As many as 50% of CD8^+^ T cells were in direct contact with LECs (Fig. 5q) and ∼65% within a 50 µm perimeter of lymphatic vessels (Fig. 5s). CD4^+^ T cells showed even closer association with lymphatics, with ∼75% in direct contact with LECs (Fig. 5p-s). NDV treatment did not affect localization of CD8^+^ T or CD4^+^ T cells relative to lymphatics (Fig. 5p-s). These data demonstrate that T cells in VEGF-C expressing tumors establish direct contact with LECs.

### Distinct changes in T-cell composition of sentinel and distant lymph nodes are associated with complete response

To assess changes in the immune composition of lymph nodes (LNs), we next analyzed tumor sentinel (sLN) and contralateral, non-sentinel lymph nodes (nsLN) by spectral flow cytometry (Fig. 6). LNs were collected 12h after the second NDV or PBS treatment of B16/EV and B16/VEGF-C tumors (Suppl. Fig. 9a). In sentinel LNs, relative frequences of total CD8^+^ T cells decreased upon NDV treatment of B16/VEGF-C tumors only, whereas total CD4^+^ T cell frequences remained unchanged across treatment groups (Fig. 6a, b, Suppl. Fig. 9b). Naïve CD8^+^ T cells (CD62^hi^CD44^low^) were markedly reduced by NDV, VEGF-C, or their combination (Fig. 6c, Suppl. Fig. 9c). Both, CM (CD62L^hi^CD44^hi^) and EM (CD62L^low^CD44^hi^) T cells were reduced only when B16/VEGF-C tumors were treated with NDV (Suppl. Fig. 9c-e). CD62L^low^CD44^low^ CD8^+^ T cells were enriched in sLNs of both, EV/NDV and VEGF-C/NDV tumors (Fig. 6d, Suppl. Fig. 9c). In contralateral non-sentinel LNs, CD8^+^ T cell subsets showed similar but less pronounced changes than in sentinel LNs following NDV treatment of B16/VEGF-C tumors; most notably, reduced total and naïve CD8^+^ T cells and increased CD62^low^CD44^low^ CD8^+^ T cells (Fig. 6a, c, d, Suppl. Fig. 9c).

**Fig 6.**
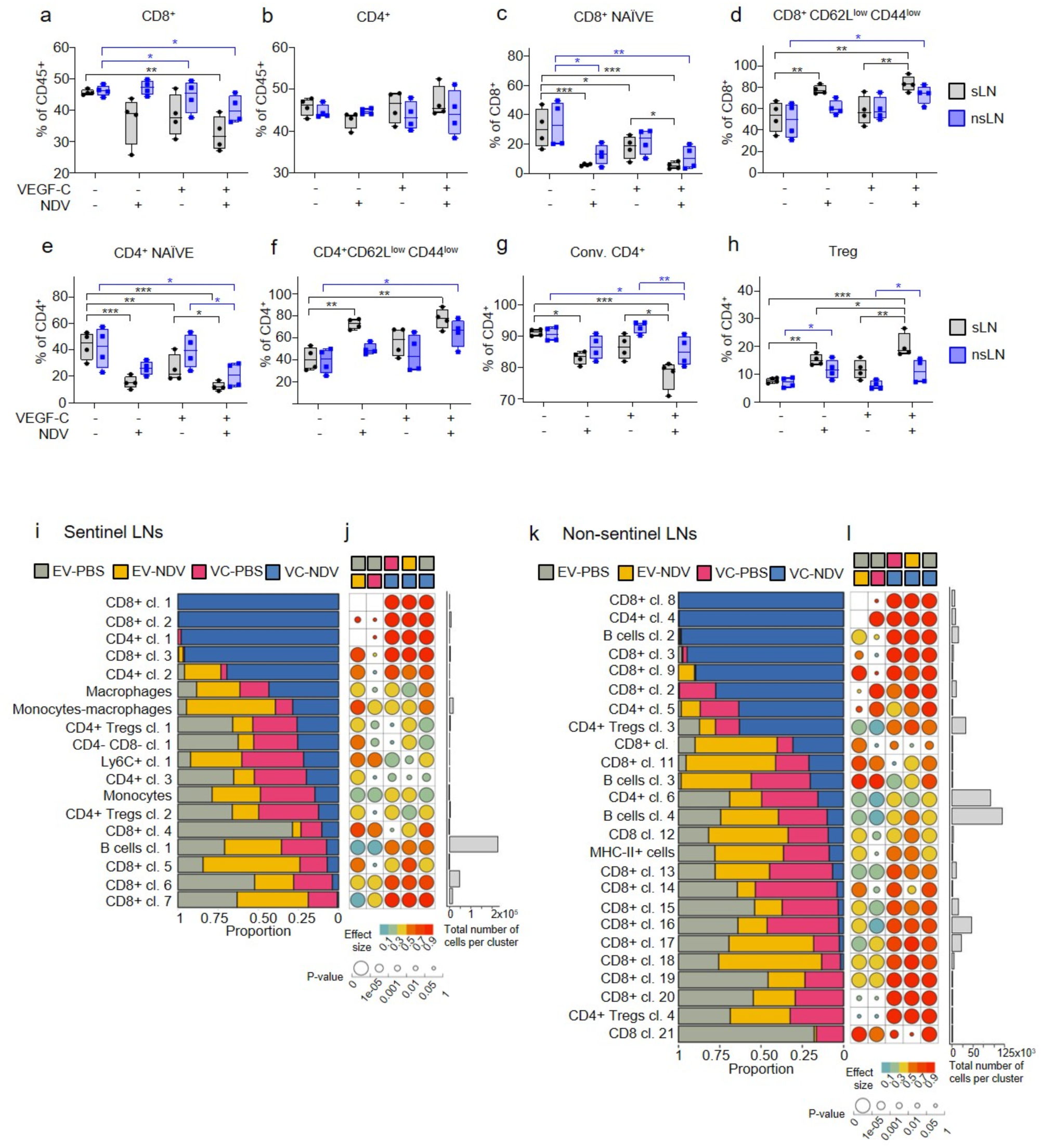
Effects of VEGF-C and NDV on immune cell composition in the lymph nodes of B16F10 tumor-bearing mice. (a-h) Spectral flow cytometry analysis of T cells in sLNs and non-sLNs. Relative frequences of distinct CD8^+^ and CD4^+^ T cell subsets are shown; total CD8^+^ (a), total CD4^+^ (b), naïve CD8^+^ T cells (c), CD8^+^ CD62L^low^ CD44^low^ T cells (d), naïve CD4^+^ T cells (e), CD4^+^ CD62L^low^CD44^low^ T cells (f), CD4^+^ Tconv (g), CD4^+^ Foxp3^+^ Tregs (h). Statistical analysis: one-way ANOVA multiple comparisons. n=4. Box-and-whiskers plots show individual values with mean ± SEM, and whiskers spanning the full range. (i,k) Stacked barplots showing cell cluster proportions in sentinel LNs (i) and non-sentinel LNs (k) across treatments. Different experimental groups are represented by their respective colors. Only cell clusters which showed significant difference between any of the conditions are shown (j, l) Heatmap shows p-values (dot size) and effect size (Cohen’s W; dot color) of the Chi-square goodness-of-fit test for pairwise comparison of cell cluster proportions across treatments. Only clusters with over 100 cells, medium or large overall effect size and a Chi-square p-value of less than 0.05 in comparisons of different treatments are depicted. The bar plot on the far right displays the total number of cells in each cluster. *p < 0.05, **p < 0.01, ***p < 0.001, not indicated = not significant.

Changes in CD4^+^ T cell subsets in sentinel LNs paralleled those of CD8^+^ T cells (Fig. 6e, Suppl. Fig. 9). Naïve CD4^+^ T cells were markedly reduced by NDV, VEGF-C, or their combination (Fig. 6e). Conversely, CD62^low^CD44^low^ CD4^+^ T cells were highly enriched upon NDV treatment, independent of VEGF-C (Fig. 6f). CD4^+^ CM and EM frequences remained unchanged (Suppl. Fig. 9 f-h). Of note, NDV triggered stark decrease in conventional CD4^+^ T cells and an increase in Tregs in sLNs, especially in combination with VEGF-C (Fig. 6g, h; Suppl. Fig. 9i). In the nsLNs, CD4 T cell changes resembled those seen in sentinel LNs but were less pronounced. Specifically, NDV/VEGF-C combination decreased frequencies of naïve and conventional CD4^+^ T cells, and increased frequencies of CD62 ^low^CD44^low^ CD4^+^ T cells and Tregs (Fig. 6e-h).

We next performed unsupervised analysis of spectral flow cytometry data from lymph nodes. NDV treatment of B16F10/VEGF-C tumors induced T cell responses in both sLNs and nsLNs, characterized by the dominant emergence of several CD8⁺ and CD4⁺ T cell clusters, along with selective depletion or marked reduction of others (Fig. 6i-l). Top three enriched CD8^+^ T cell clusters were all positive for Ly6C and likely represent recently activated effector CD8^+^ T cells (Cl. 1 and 3, CD44^+^Ly6C^+^; Cl. 2, CD103^+^Ly6C^+^). This response was particularly striking in nsLNs, where multiple CD8⁺ and CD4⁺ T cell subsets selectively expanded, while several others were markedly reduced. These data demonstrate that NDV and VEGF-C induce changes in both sentinel and in distant LNs, suggesting the importance of systemic effects in achieving tumor elimination.

### VEGF-C is widely expressed in human cancers

We next evaluated VEGF-C mRNA expression across 31 human cancer types by analyzing bulk RNA-seq data from the TCGA Pan-Cancer Atlas (PANCAN) (Fig. 7). VEGF-C was expressed in all solid tumor types analyzed, and it was highly expressed in 47.7% of all tumor samples in the database, where high expression is defined as log₂(TPMK) ≥ 7, corresponding to the median value across all tumors (Fig. 7a). High VEGF-C expression was observed in 13 tumor types, including bladder cancer (BLCA), breast cancer (BRCA), head and neck cancer (HNSC), kidney cancer (KICH, KIRC), lung adenocarcinoma (LUAD), mesothelioma (MESO), pancreatic adenocarcinoma (PAAD), neuroendocrine tumors (PCPG), sarcoma (SARC), and thyroid cancer (THCA) (Fig. 7a). Notably, 100% of thyroid cancers (THCA), 80% of sarcomas (SARC), 79% of pancreatic cancers (PAAD) and 65% of head and neck cancers (HNSC) exhibited high VEGF-C expression (Fig. 7b). In contrast, glioblastoma multiforme (GBM), low-grade glioma (LGG), and adenocarcinomas of the colon and rectum (COAD, READ), were among the lowest VEGF-C-expressing tumors (Fig. 7a, b). In cutaneous and uveal melanomas (SKCM, UVM) VEGF-C expression was ubiquitous with intermediate expression levels. These findings highlight a subset of tumors with high VEGF-C expression, suggesting that they may be more amenable to APMV-based virotherapy.

**Fig 7.**
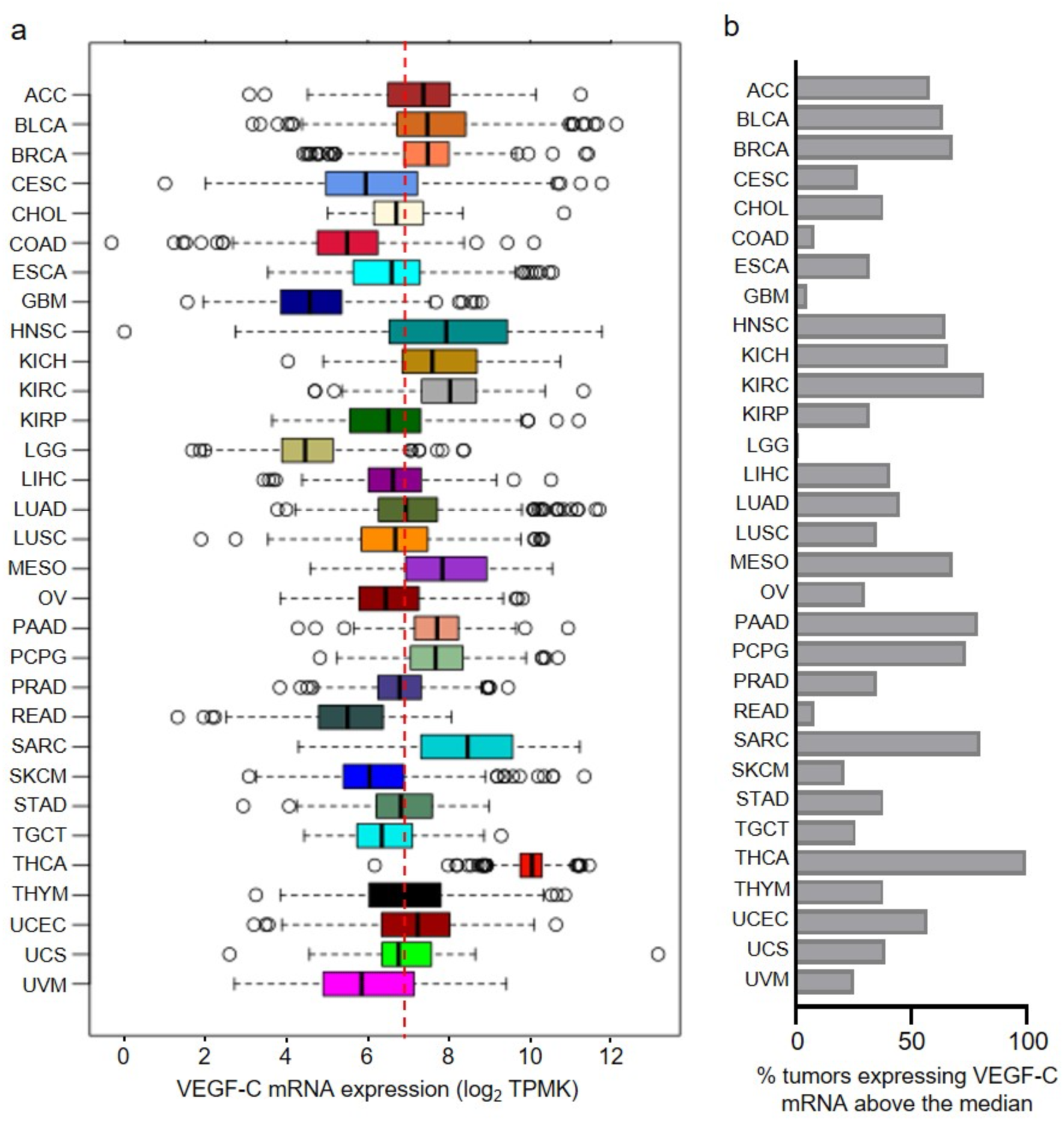
VEGF-C mRNA expression in human tumors. (a) Gene expression data from the TCGA PANCAN 2022 cohort showing expression of VEGF-C in human solid tumors (n=10,839). The red dotted line indicates median VEGF-C expression across all solid tumors in the cohort. (b) Percentage of tumors within each cancer type expressing VEGF-C above the pan-cancer median. Tumor type abbreviations follow The Cancer Genome Atlas (TCGA) nomenclature: ACC (Adrenocortical carcinoma), BLCA (Bladder urothelial carcinoma), BRCA (Breast invasive carcinoma), CESC (Cervical squamous cell carcinoma and endocervical adenocarcinoma), CHOL (Cholangiocarcinoma), COAD (Colon adenocarcinoma), ESCA (Esophageal carcinoma), GBM (Glioblastoma multiforme), HNSC (Head and neck squamous cell carcinoma), KICH (Kidney chromophobe), KIRC (Kidney renal clear cell carcinoma), KIRP (Kidney renal papillary cell carcinoma), LGG (Low-grade glioma), LIHC (Liver hepatocellular carcinoma), LUAD (Lung adenocarcinoma), LUSC (Lung squamous cell carcinoma), MESO (Mesothelioma), OV (Ovarian serous cystadenocarcinoma), PAAD (Pancreatic adenocarcinoma), PCPG (Pheochromocytoma and paraganglioma), PRAD (Prostate adenocarcinoma), READ (Rectum adenocarcinoma), SARC (Sarcoma), SKCM (Skin cutaneous melanoma), STAD (Stomach adenocarcinoma), TGCT (Testicular germ cell tumors), THCA (Thyroid carcinoma), THYM (Thymoma), UCEC (Endometrial cancer), UCS (Uterine carcinosarcoma), and UVM (Uveal melanoma).

## DISCUSSION

Our results demonstrate exceptional anti-tumor efficacy of APMV-based viral immunotherapies in presence of VEGF-C in tumors. Treatment of B16F10 tumors with either NDV or APMV-4 oncolytic viruses led to complete remission in most animals when tumors expressed VEGF-C. Responses were durable, with long-term survival, protection from tumor development upon re-challenge, and absence of metastases, indicating the establishment of robust immunological memory. These results identify a combination of APMV and VEGF-C as a highly effective anti-tumor therapeutic approach leading to durable responses and highlight a novel strategy for improving efficacy of oncolytic viruses by activating the lymphatic system.

It is important to emphasize that we show exceptional anti-tumor efficacy of APMVs as a monotherapy. OVs generally show limited efficacy as a single agent and primarily delay tumor progression(*11, 15, 40*). NDV, for example, has been shown to impair tumor growth and prolong survival across multiple studies, but it rarely leads to tumor eradication(*4–6, 15, 36, 40*). Most current strategies for improving efficacy of OVs focus on enhancing anti-tumor immune responses by targeting immune cells directly. These approaches typically involve engineering viruses to express immunostimulatory cytokines, superagonists for co-stimulation, or combining OVs with systemic immunotherapies such as anti-PD-1 or anti-CTLA-4(*4–7, 21, 41–43*) and often require concurrent delivery of multiple immunomodulators to achieve robust therapeutic outcomes. For example, NDV expressing IL-12, CD28 superagonist, and systemic PD-1 blockade achieved 37% complete response in a mouse model(*21*), whereas achieving complete response rates greater than 70% required combining NDV concurrently with blocking antibodies against PD-1, PD-L1, and CTLA-4(*18, 40*). NDV treatment with anti-PD-1 alone led to only 20% CR in B16F10 model(*4, 5*).

In striking contrast, we observed complete response (CR) rates exceeding 90% using NDV or APMV-4 as single-agent therapy in VEGF-C-expressing tumors, without any additional immunomodulatory agents. These findings underscore the central role of VEGF-C and lymphatics in orchestrating anti-tumor immunity. Seminal work by Fankhauser and Swartz (2017) first demonstrated that VEGF-C-mediated lymphangiogenesis can enhance immunotherapy efficacy in preclinical models of melanoma(*34*). Several immunotherapy strategies including adoptive T cell transfer, DC vaccination, and CpG adjuvant have shown prolonged survival and 10-20% CR using B16F10/VEGF-C tumors, but only in the presence of a potent model antigen(*34*). Cancer vaccine using irradiated tumor cells engineered to secrete VEGF-C (VEGF-C vax) conferred prophylactic protection against melanoma and improved responses when combined with immune checkpoint blockade, but did not induce complete remission(*35*). Similarly, prophylactic delivery of AAV-VEGF-C protected against glioblastoma (GBM) development(*44*). In therapeutic setting however, PD-1 blockade combined with VEGF-C achieved high CR rates only when GBM tumor burden was low, while in established tumors, dual PD-1 and TIM-3 blockade was necessary to extend survival but did not induce CR(*44*).

A key distinction of our study is the exceptionally high survival benefit achieved with VEGF-C and APMV alone in an intervention treatment setting. Furthermore, we demonstrate long-term survival without tumor recurrence beyond two years, whereas previous studies had endpoints at 50-100 days(*34, 35, 44*). Notably, even after this extended period of observation, no metastases were detected. This finding is particularly significant given the well-established role of tumor-derived VEGF-C in promoting metastatic spread(*45, 46*). Our results underscore the fundamentally different outcome of VEGF-C expression in the context of immune activation, where it orchestrates immune response and contributes to durable tumor control and suppression of metastases.

The combination of VEGF-C and APMV reprograms the tumor immune microenvironment in multiple ways to achieve tumor regression and we identified several hallmark changes associated with complete response: (1) increased infiltration of CD8⁺ and CD4⁺ T cells; (2) activation of T cells and NK cells; (3) expansion of specific effector and memory T cell subsets; and (4) reduction of inhibitory CD4⁺ T cells and immunosuppressive cues.

In the immunologically cold B16F10 melanoma model, VEGF-C expression induced lymphangiogenesis and markedly increased intratumoral densities of CD8⁺ and CD4⁺ T cells. This is consistent with previous studies linking lymphatic expansion to T cell accumulation(*34, 47*). T cell infiltration is a well-established predictor of patient survival and response to immunotherapy across multiple cancer types(*48–52*). In our study increased T cell presence alone was not sufficient to elicit anti-tumor responses, but it was a prerequisite for mounting effective anti-tumor immunity upon APMV treatment. Consistent with this, VEGF-C-expressing tumors harbored few CD8⁺ and CD4⁺ effector T cells, a low CD8⁺ effector-to-Treg ratio, and a high proportion of CXCR3+TIM3+ early exhausted CD8⁺ T cells. These findings align with prior reports indicating that VEGF-C alone does not impair tumor growth but may contribute to the establishment of a tolerogenic tumor microenvironment(*53*).

APMV treatment of tumors in which TME has been altered with VEGF-C, shifts the tumor immune landscape toward effector T cell phenotypes. Some of these changes were driven independently by VEGF-C or APMV, and their combination likely produced additive effects, while some emerged only with the VEGF-C/APMV combination, indicating synergy. One such synergistic effect was the unique expansion of CD8^+^CD44^+^PD-1^+^CD25^+^ T cells correlated with CR. These cells represent recently activated T cells in our model, since they were detected only 2.5 days after treatment initiation.

Prior studies have shown that PD-1 upregulation is a hallmark of early antigen-specific T cell activation (*54*) and that the expression of co-inhibitory receptors alone does not indicate dysfunctional T cell phenotype in melanoma (*55, 56*). CD25^+^PD-1^+^CD8^+^ T cells have been identified as a key mediators of effective antitumor immunity (*57, 58*) and they serve as a marker of effective T cell activation following DC vaccination in melanoma (*59, 60*). In our study, robust expansion of CD25^+^PD-1^+^ effector CD8^+^ T cells and reduction of Tregs in NDV-treated B16/VEGF-C tumors, led to increased ratio of CD25^+^PD-1^+^CD8^+^ T cells to Tregs that was indicative of CR.

The mechanisms by which VEGF-C and APMV drive the expansion of CD8^+^CD25^+^PD-1^+^ T cells remain to be elucidated. VEGF-C-mediated lymphangiogenesis may create a supportive niche for proliferation or retention of intratumoral TCF-1+ stem-like CD8+ T cells, which serve as precursors to CD8^+^PD-1^+^ effector cells and are essential for sustaining effector responses (*61*). Notably, both TCF-1^+^ and memory T cells have been shown to egress to sentinel LNs via lymphatics and are continuously replenished in tumors by newly recruited cells (*62, 63*). Thus, expansion of lymphatic vasculature may facilitate recirculation of T cells, thereby limiting exhaustion and supporting the maintenance of functional effector populations.

Furthermore, APMV in combination with VEGF-C rapidly activates multiple effector cell populations, including NK cells, CD8^+^ and CD4^+^ T cells. APMV alone strongly activated NK cells and induced GzmB+, in agreement with prior studies (*6*). However, in the presence of VEGF-C, APMV increased GzmB+ CD8+ T cells, polyfunctional CD8 T cells and CD4+ effector T cells, while reducing exhausted-like TIM-3+ CD8+ subsets (*55, 64*), thus further increasing overall cytotoxic potential. While VEGF-C alone expanded several effector-like CD8^+^ subsets, APMV was necessary to trigger cytokine and GzmB production, emphasizing that VEGF-C creates a receptive immune landscape, but APMV is required to trigger effector engagement. The unique efficacy of APMV and VEGF-C combination likely reflects synergy of NK and T cell responses, where APMV alone potently stimulates NK cells, and in combination with VEGF-C also potently enhances T cell activation. Unlike checkpoint blockade, which primarily targets adaptive immunity, this approach potently engages both innate and adaptive arms of the immune system.

In addition to promoting effector responses, VEGF-C/APMV treatment reduced immunosuppressive CD4⁺ T cell subsets expressing CD25 and CTLA-4, likely antigen-experienced cells which consume IL-2 and dampen CD8⁺ T cell responses. Their diminished presence may contribute to reduced suppression, enabling sustained CD8⁺ T cell cytotoxicity. It has been suggested that one reason that NDV as a single agent may not be sufficient to drive complete tumor rejection is because of inhibitory mechanisms induced in response to the virus subsequent to activation (*40*). In line with this concept, we show that while NDV initiated effector activation, it also increased suppressive CD4^+^ T cells that may limit durable responses. VEGF-C countered this feedback inhibition by reducing frequencies of CD4^+^CTLA-4^+^ Tregs, thus enabling sustained co-stimulation and cytokine support. Hence, VEGF-C reduces inhibitory signals and effectively abolishes CTLA-4-mediated suppression.

We demonstrate here that VEGF-C mRNA expression is a common feature across many solid tumors. Our findings suggest that high VEGF-C expression may sensitize tumors to APMV-based virotherapy, potentially identifying a subset of patients most likely to benefit clinically. For tumors with low endogenous VEGF-C expression, exogenous delivery of VEGF-C may enhance therapeutic efficacy of OVs. In melanoma, our data suggest that patients with high VEGF-C expression may be more likely to respond favorably. In support of this idea, a clinical trial has shown that melanoma patients with elevated serum VEGF-C levels experienced increased progression-free survival following checkpoint inhibitor therapy (*34*). Further studies are needed to determine whether VEGF-C could help predict therapeutic responses to APMVs and other oncolytic viruses.

In conclusion, in presence of VEGF-C, APMVs achieve complete response through multipronged mechanisms; by increasing immune cell infiltration in tumors and by enhancing, accelerating and sustaining immune activation. Our findings mark a significant advance by demonstrating the curative potential of APMVs when VEGF-C is expressed in tumors. This lymphatic-focused approach represents a promising novel strategy for enhancing efficacy of immunotherapies without overlapping toxicities, particularly in tumors where checkpoint blockade has had limited impact.

## MATERIALS AND METHODS

### Cells and culture conditions

B16F10 cells were culture in DMEM (Gibco, cat. 11960-044) supplement with 10% FBS (Corning 35-010-CV), L-glutamine 200mM (Gibco, cat. 25030-81) and Antibiotic-Antimycotic solution (Gibco, cat. 15240-062). Vero african green monkey kidney epithelial cells (ATCC Cat# CCL-81) were maintained in DMEM medium supplemented with 10% FBS, 1% Glutamax-100X (Invitrogen), penicillin and streptomycin (*1*). We received B16F10 – VEGF-C cells from Dr. Melody Swartz, (The University of Chicago, Illinois, USA). To generate B16F10 empty vector (EV) cells we produced lentiviruses using CaCl_2_ transfection protocol. These lentiviruses carried a mammalian-expression empty vector (pD2109, ATUMbio) (*35*). Briefly, viral envelope-encoding plasmids (pMD.2 and psPAX2) were mixed with empty vector (EV) plasmid in a CaCl_2_-HBSS 2X solution. Transfection particles were allowed to form for 30 minutes at room temperature and 1 ml of the solution was applied drop wise to an 80% confluent 293t 10 cm dish. 24 hours after transfection the conditioned media containing viruses was collected, centrifuged at 1200 rpm for 5 minutes and filter through a 45 µm filter, aliquoted and stored at −80°C. Then, 2×10^5^ B16F10 cells were seeded in a 60 mm dish supplemented with culture media. When cells reached 50% confluency were transduced with 1 ml of the previously described lentivirus containing media plus 1 ml of culture media. After 48 hours the media was replaced, and cells were grown for another 72 hours to allow cells to express the desired transgene and the puromycin resistance gene. When cells were confluent, they were transferred to a 10 cm dish. After 72h, media was replaced with selection media containing 2 µg/ml puromycin (Sigma, cat. 540411). Cells were then culture in selection media for 2 weeks.

### Avian Paramyxoviruses

Modified NDV virus LaSota-L289A has been previously described (*65*). APMV-4 Duck/Hong Kong/D3/1975 infectious clone used in these studies was previously reported (*36*). Viral stocks were propagated in 9-day-old embryonated chicken eggs and clear purified from the allantoic fluid by discontinuous sucrose density gradient ultracentrifugation for resuspension and storage in PBS. Viral titers were calculated by indirect immunofluorescence on Vero cells using serotype-specific antiserums to NDV (Rabbit polyclonal(*66*)) and to APMV-4 (475-ADV chicken polyclonal antiserum; USDA/NVSL). Secondary antibodies, goat anti-rabbit Alexa Fluor 488 (A-11008) and anti-chicken Alexa Fluor 488 (A-11039) were purchased from Invitrogen.

### Western Blot

An 80% confluent 10 cm dish culture media was replaced with 1% FBS serum media for each cell line to be analyzed for 12h. After 12h, media was collected, centrifuged at 1200 rpm for 5 minutes at 4°C, filtered through a 40 µm filter (Fisher Scientific, cat. 22-363-547), aliquoted and stored at −80°C. 8 ml of the media were then concentrated using Amicon 3K columns for protein concentration (Sigma Millipore cat. UFC900324) for 1h at 4°C at 4000g. The collected supernatant was equalized to 300 µl with serum free media. 15 µl of the concentrated media were supplemented with 5 µl of 6x Laemmli buffer, boiled at 96°C for 5 minutes and loaded in a 15% acrylamide SDS-PAGE gel. The resolved gel was transferred to a PVDF membrane (Biorad, cat. 1620177). Primary anti-VEGFC antibody (R&D, AF752) was diluted 1/100 in 5% milk-PBS-Tween 20 0.2% and incubated RT for 1h. Secondary anti-goat HRP (Jackson Immunoresearch, Inc., Cat. 705-036-147) antibody was diluted 1:200 in 5% milk-PBS-Tween 20 0.2% and incubated RT for 1h.

### RNA Isolation and qPCR

RNA was extracted using QIAGEN RNeasy mini KIT (QIAGEN, cat.74104) according to manufacturer instructions from a 90 – 100% confluent 10 cm dish of each cell line. 1 µg of RNA was used to produce cDNA using iScript™ cDNA Synthesis Kit (Biorad, cat. 1708890) according to manufacturer instructions. Resulting cDNA was diluted 1/10 and 2 µl of the dilution were used per reaction. qPCR was run in a 96 well plate using PowerUp™ SYBR™ Green Master Mix (Applied Biosystems, A25741) according to manufacturer instructions in a OneStep qPCR machine (Applied Biosystems). Transcript expression was normalized to the Actin and GADPH house-keeping gene and mRNA[RU] was calculated using the 2^-ΔΔCT^ method for qPCR analysis. Primer sequences are described in the detailed methods (Extended Methods Table 4).

### ELISA

An 80% confluent 10 cm dish culture media was replaced with 1% FBS serum media for each cell line to be analyzed for 12h. After 12h, media was collected, centrifuged at 1200 rpm for 5 minutes at 4°C, filtered through a 45 µm filter, aliquoted and stored at −80°C. 100 µl of the conditioned media were used according to manufacturer instructions (R&D, cat. DVEC00) to measure VEGF-C expression through ELISA. When the conditioned media was collected cells were counted for normalization.

### Mouse experiments and tissue collection

B16F10-EV or B16F10-VEGF-C tumor cells (3 x 10^5^ cells in 100 µl serum-free media) were injected intradermally into the right flank of seven to ten-week-old C57Bl/6J mice (Jackson, cat.000664). Mouse weights and tumor sizes were measured every two days. Treatment was started when tumors reached 5 mm in size. 50 µL of a solution containing PBS, NDV or APMV-4 (10^7^ PFUs/dose in 50 µL) were administered to the mice intratumorally every two days for a total of 4 treatments. Mice were monitored until humane endpoint. Mouse experiments were performed in accordance with protocols approved by the Institutional Animal Care and Use Committee (IACUC). The formula 4/3π(length/2)^2^(width/2) was used to calculate tumor volume.

### Immunofluorescent staining and quantification

For cell culture immunofluorescence, a well of a glass slide (ICN Biomedical, Cat.6041805) was seeded with 5×10^4^ cells. After 12h, cells were washed with PBS and fixed in 10% formalin for 10 minutes RT. Fixed cells were blocked with 1% goat serum in PBS-Tween 20 0.2% for 1h RT. Cells were then incubated with anti-VEGFC primary antibody (R&D, cat. AF752) diluted 1/100 in 1% goat serum-PBS-Tween 20 0.2% and incubated RT for 1h. Secondary anti-goat Alexa-594 antibody (Jackson Immunoresearch, Inc, cat. 705-586-147) was diluted 1:500 in 1% goat serum PBS-Tween 20 0.2% and incubated RT for 1h. Nuclei were counterstained with DAPI (1 μg/ml). Slides were coverslipped using fluorescence mounting medium (Dako cat. S3023). Images were captured using the Nikon Eclipse E600 microscope and DS-Qi1 monochrome camera. For tissue immunofluorescence, fresh mouse tissues were embedded in OCT (ThermoFisher, cat. 23-730-571) and sectioned at 6-7 µm on Cryostat (Leica Biosystems, CM3050S). Sections were either stained immediately or stored at −80°C. Sections were fixed in acetone on ice for 2 minutes, followed by 80% methanol for 5 minutes. Sections were washed with PBS (Dulbecco’s, no Ca or Mg, Gibco, cat. 14-190-250) 3 times for 5 minutes. Sections were then incubated in primary antibody (Extended Methods Table 2) in antibody diluent (PBS, 12% BSA, 0.01% NaN3) in a humidified chamber. Sections were washed 3 times for 5 minutes in PBS and then incubated in the corresponding fluorescent conjugated secondary antibody for 1 hour at room temperature in a dark humidified chamber. Sections were washed again 3 times for 5 minutes in PBS and incubated in DAPI 1 µg/mL for 30 seconds. Sections were washed 3 times for 5 minutes in PBS and coverslipped using DAKO Fluorescent Mounting Media (Agilent, S3023).

### Immunohistochemistry

Immunohistochemistry was performed on paraffin-embedded tissue sections using the Leica Bond RX automated Immunostainer (Leica Biosystems), according to the Leica staining protocol. Briefly, all slides were deparaffinized using a heated Bond™ Dewax Solution (Leica cat. AR9222) and washed with 1X Bond™ Wash Solution (Leica cat. AR9590). Epitope retrievals were carried out for 20 minutes using the citrate-based Bond™ Epitope Retrieval 1 solution (Leica cat. AR9961) or solution 2 (Leica cat. AR9961). Slides were then blocked with either the Dako Dual Endogenous Enzyme Block (Agilent, S2003) for ten minutes prior to staining with the Bond™ Polymer Refine Red Detection System (Leica cat. DS9390), or with a 3-4% v/v hydrogen peroxide block for five minutes that is incorporated into the Bond™ Polymer Refine Detection kit (Leica cat. DS9800) and used for DAB staining. Melan A antibody was diluted in Bond™ Primary Antibody Diluent (Leica Biosystems, AR9352). Slides were mounted Glycergel Aqueous Mounting Media (DAKO, cat. C0563) or Cytoseal XYL (Fisher Scientific, cat.50-209-0573). A list of antibodies and experimental conditions can be found in Extended Methods Table 3 and 4.

### Spectral Flow Cytometry

Spectral Flow cytometry was performed using Aurora Spectral Cytometer (Cytek Biosciences) or and flow cytometry using LSR Fortessa X-20 (BD Biosciences). In brief, tissues were dissected and minced in a sterile petri dish in ice cold PBS. Tumor tissues were dissociated with Mouse Tumor Dissociation Kit (Miltenyi, cat. 130-096-730) enzymes in Octomacs Dissociator with Heaters (Miltenyi, cat. 130-096-427). Lymph nodes were dissociated with Collagenase D enzyme (1 mg/ml, Roche, cat. 50-100-3282) in 37°C water bath for 1h. Dissociation reactions were stopped with the addition of ice-cold FACS buffer (1%FBS, 0.09% NaN3 in PBS). Erythrocytes were lysed using RBC lysis (cat. 00-4300-54, eBioscience) for 1 minute on ice. Lysis was stopped by the addition of ice-cold FACS buffer. Dissociated tissues were pressed through a 70 µm nylon filter (Fisher Scientific, cat. 22-363-548) to create a single cell suspension. Cell yield and viability were determined using Countess II Automated Cell Counter (ThermoFisher). Samples were stained with primary antibodies (Extended Methods Table 1) targeting cell surface markers for 30 minutes on ice (10^6^ cells/100 µL). They were then fixed and permeabilized with FOXP3 Transcription Factor Staining Buffer Set (cat. <u>00-5523-00</u>, eBioscience). Samples were then stained with primary antibodies targeting intracellular markers. Compensation and reference groups were calculated using UltraComp beads (cat. 01-2222-41, eBioscience). Dead cells were excluded using LIVE/DEAD Fixable Yellow Dead Cell Stain Kit (cat. L34968, Invitrogen). 2.0×10^4^-1.0×10^5^ viable CD45^+^ cells were acquired/sample. Flow cytometry data analysis was performed on FCS Express Version 7.0.

### Image processing and analysis

Slides were scanned with Nanazoomer Digital Slide Scanner (Hamamtsu, Shizuoka, Japan). Image processing was performed using NDP viewer 2 (Hamamatsu, Shizuoka, Japan) and analyzed using Fiji open-source software v.1.53q (National Institutes of Health). Confocal images were acquired at the Microscopy and Advanced Bioimaging CoRE at the Icahn School of Medicine at Mount Sinai using a ZEISS LSM 780 Laser Scanning Microscope (ZEISS, USA). Image processing and analysis was performed using Fiji open-source softwarev.1.53q (National Institutes of Health). For quantification of cells, single channel 8-bit scanned images of complete tumor sections were analyzed using count particles Fiji tool. 3 to 5 different whole tumor sections were counted per experimental group. T cell distance to LYVE-1^+^ cells was performed using Qupath (v 0.5.1). Regions of interest were drawn for each tumor we done using the polygon annotation tool. Next, cell detections were performed within each region of interest using DAPI (blue) channel to detect the nucleus of individual cells. LYVE-1 (green) and CD8 or CD4 (red) channels were used to detect and quantify LYVE-1^+^, CD8^+^ and CD4^+^ cells, respectively. Spatial analysis was performed to detect centroid distances in 2D and distance annotations in 2D.

### Aurora spectral flow cytometry data analysis

High-dimensional flow cytometry data was clustered using Clustergrammer web-based tool (*67*). To ensure that each treatment group is represented by the same number of cells, we randomly subsampled cells from mice subjected to the same treatment. The number of subsampled cells per treatment was equal to the lowest number of cells over all treatments. The proportions of cells subsampled from each mouse were held constant to reflect the original proportions over all mice belonging to one treatment. Using the subsampled data set we tested whether there is a significant difference in the distribution of cells coming from different treatments using the Chi-square test for overall difference in the distribution of phenotypes across treatments and for pairwise comparisons of all combinations of different treatments. P-values were corrected for multiple testing using the Bonferroni correction. Effect size of the Chi-square test was evaluated by Cohen’s W (reference) for overall and pairwise differences in distribution Cohen’s W was calculated using the implementation in the Rcompanion package for R statistical environment (Mangiafico, 2022). We considered Cohen’s W larger or equal to 0.5 to represent a large effect size, 0.30 – 0.50 medium effect size, 0.1 – 0.3 small effect and less than 0.1 no effect. We removed all the phenotypes where the cell type could not be precisely determined and for the remaining phenotypes, we calculated the median of marker expression values (signal intensity) for clusters of cells belonging to each phenotype. 9 populations in the non-draining lymph node with the same markers and same distribution were condensed into 4 different populations to form B cells 2, CD8^+^ T cells 2, CD8^+^ T cells 14, and CD8^+^ T cells 15. The observed proportion, relative effect size and p-value of these population was not significantly altered after the merge.

### TCGA analysis

TCGA Pan-Cancer Atlas (PANCAN) 2022 transcriptomic data sets (bulk RNA-seq) were analyzed using hegemon web-based tool (http://hegemon.ucsd.edu/Tools/). VEGF-C mRNA expression was analyzed across 31 solid tumor types, comprising a total of 10,839 tumor samples. Tumors were classified as having high or low VEGF-C expression based on the median log₂(TPMK) value across all tumors from different cancer types.

### Statistical analyses

Statistical analyses were performed using GraphPad Prism (v.10) software and each specific statistical test is indicated in the corresponding figure legend.

## List of Supplementary Materials

Supplementary Materials and Methods

Supplementary Figures S1 to S9

Data files

Extended methods tables

## ACKNOWLEDGEMENTS

We would like to thank the expertise and assistance of the Dean’s Flow Cytometry CORE and the Microscopy and Advanced Bioimaging CoRE at the Icahn School of Medicine at Mount Sinai. This research was supported in part by the Tisch Cancer Institute at Mount Sinai P30 CA196521 – Cancer Center Support Grant.

## Author contributions

Conceptualization: RFR, SCC, AGS, MS

Methodology: RFR, SCC, AE, AR, NDF, IM, RK, AH

Investigation: RFR, SCC, AE, AR, YB, RK, IM, AK, AH

Visualization: RFR, RK

Funding acquisition: AGS, MS

Project administration: RFR, SCC, AGS, MS

Supervision: MS, AGS

Writing – original draft: RFR, MS

Writing – review & editing: RFR, SCC, YB, AK, AH, AGS, MS

