## Supplementary material for "Lymphangiogenesis driven by VEGF-C reshapes tumor immune landscape and enables tumor eradication by viral immunotherapy": Supp. Figs. 1-9

### Suppl. Figure 1

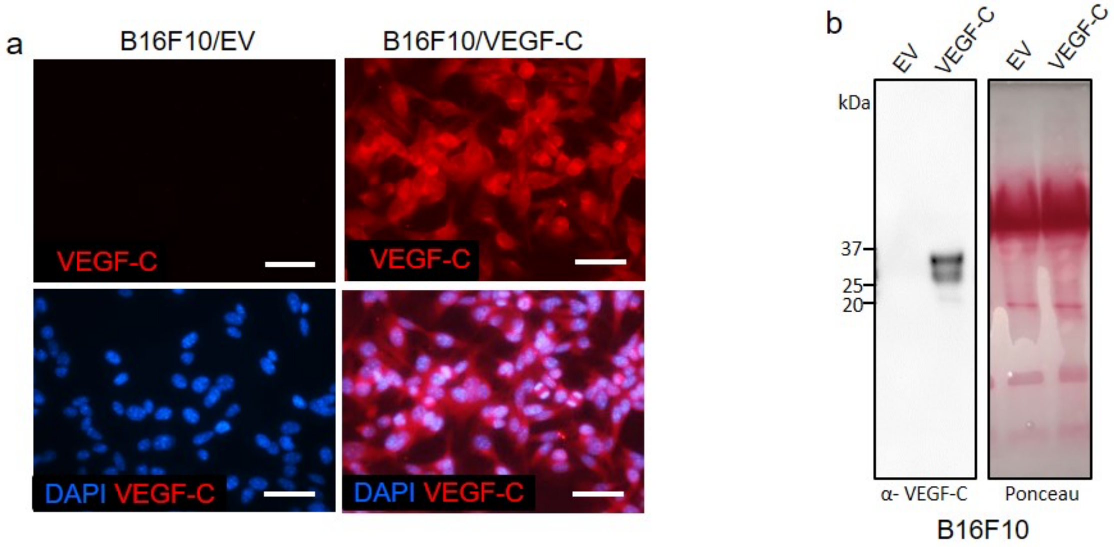


**Suppl. Fig 1. VEGF-C protein expression by B16F10 cells.** (a, b) VEGF-C expression by B16F10/EV and mVEGF-C-transduced cells analyzed by (a) immunostaining, and (b) Western blot of cells in vitro. Scale bars = 100 µm

### Suppl. Figure 2

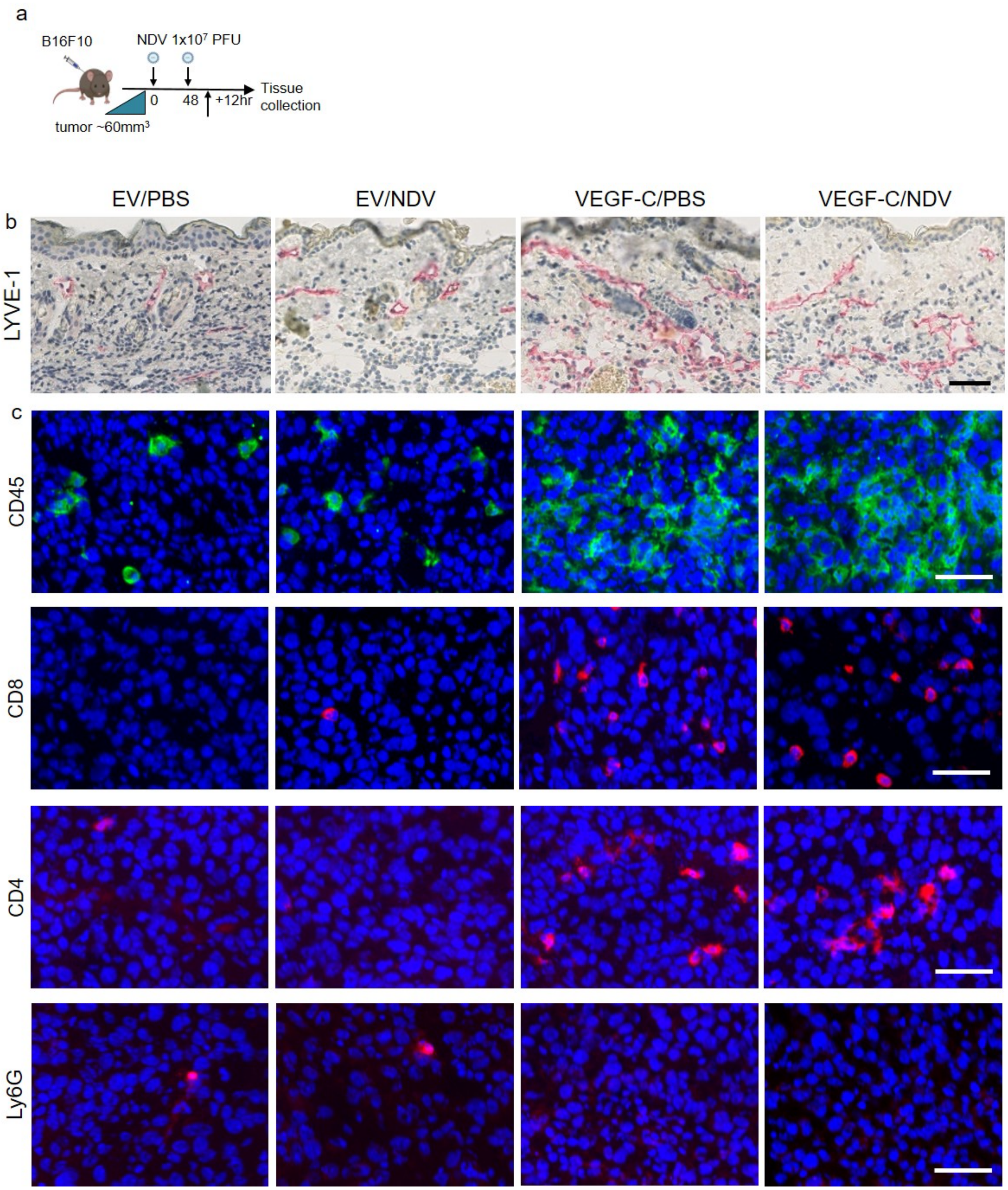

**Suppl. Fig 2. Effects of NDV and VEGF-C on lymphangiogenesis and T cell densities.** (a) Experimental design for sample collection: mice were injected with B16/EV or B16/VEGF-C cells and treated with NDV or PBS when tumors reached ~60mm$^3$. Samples were collected 12h after the second treatment. (b) Lymphatics in the skin overlying B16 tumors immunostained for LYVE-1 by IHC. (c) Immunofluorescent staining of B16 tumors for CD45, CD8, CD4, and Ly6G across treatments. Scale bars: 50 µm.

#### Suppl. Figure 3

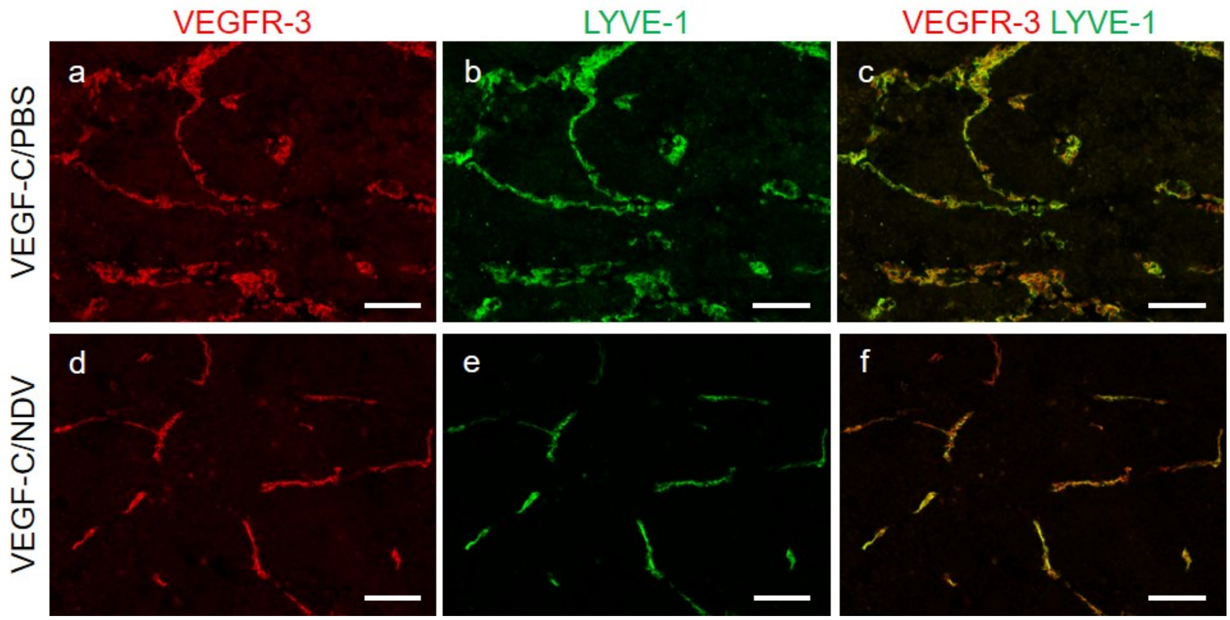


**Suppl. Fig 3. VEGFR-3 expression in B16/VEGF-C tumors.** Immunostaining for VEGFR-3 (red) (a, d), LYVE-1 (green) (b, e) and co-localization of VEGFR-3 and LYVE-1 (yellow) (c, f) in B16/VEGF-C tumors treated with PBS or NDV, as indicated. Scale bars: 100 µm

### Suppl. Figure 4

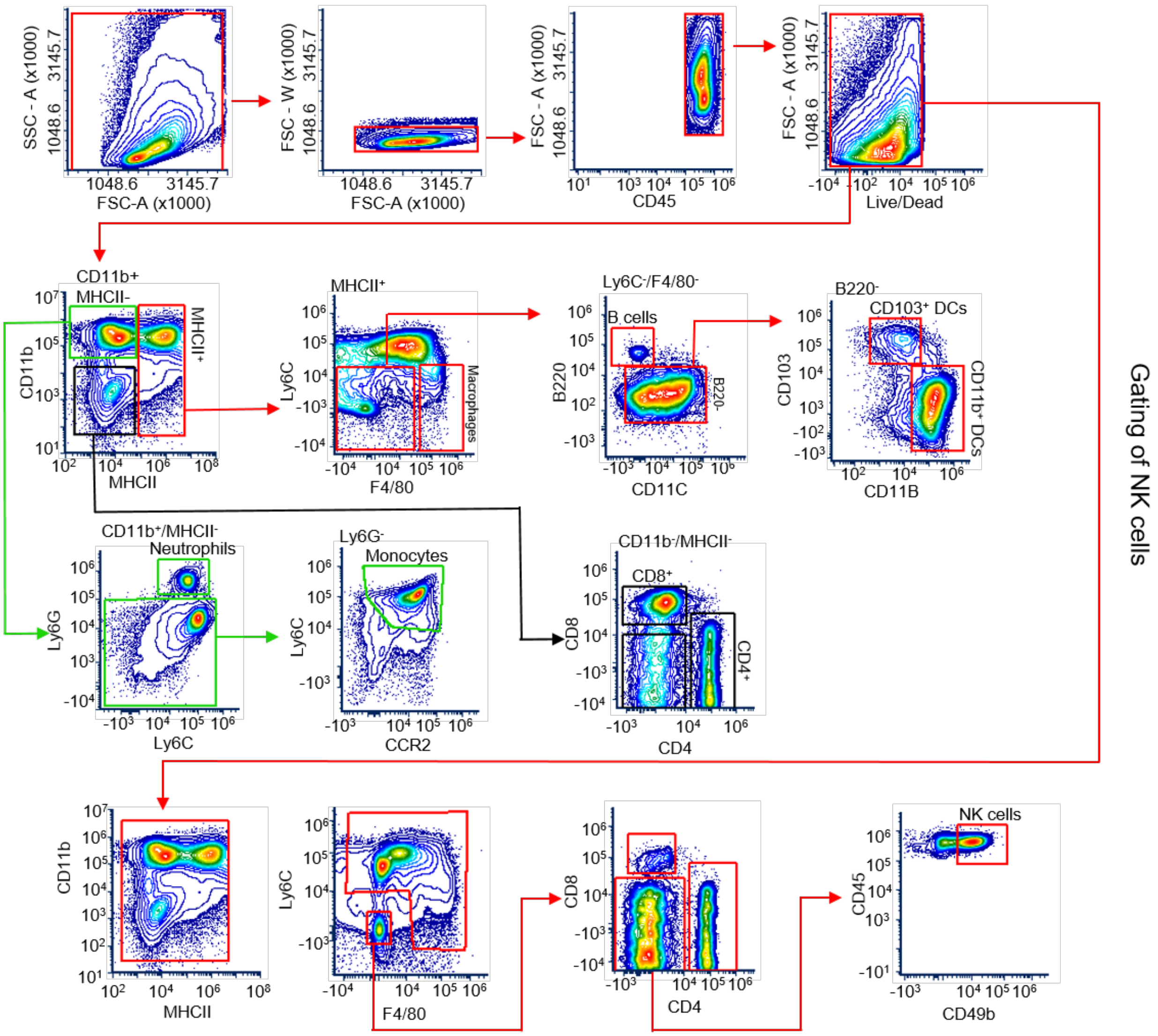


| Lineage markers | |
|---|---|
| CD4 | CD45$^+$ MHCII$^-$ CD11b$^-$ CD4$^+$ |
| CD8 | CD45$^+$ MHCII$^-$ CD11b$^-$ CD8$^+$ |
| Macrophages | CD45$^+$ MHCII$^+$ Ly6C$^-$ F4/80$^+$ |
| Monocytes | CD45$^+$ MHCII$^-$ CD11b$^+$ Ly6G$^-$ Ly6C$^+$ |
| Neutrophiles | CD45$^+$ MHCII$^-$ CD11b$^+$ Ly6G$^+$ Ly6C$^{low}$ |
| B cells | CD45$^+$ MHCII$^+$ Ly6C$^-$ F4/80$^-$ CD11c$^-$ B220$^+$ |
| NK cells | CD45$^+$ Ly6C$^-$ F4/80$^-$ CD4$^-$ CD8$^-$ CD49b$^+$ |
| Total DCs | CD45$^+$ MHCII$^+$ Ly6C$^-$ F4/80$^-$ CD11c$^-$ B220$^-$ |
| CD103$^+$ DCs | CD45$^+$ MHCII$^+$ Ly6C$^-$ F4/80$^-$ CD11c$^-$ B220$^-$ CD103$^+$ CD11b$^-$ |
| CD11b$^+$ DCs | CD45$^+$ MHCII$^+$ Ly6C$^-$ F4/80$^-$ CD11c$^-$ B220$^-$ CD103$^-$ CD11b$^+$ |

**Suppl. Fig 4. Flow cytometry gating strategy for main immune cell subtypes.** Manual gating scheme for delineating subsets of myeloid and lymphoid cells analyzed from the spectral flow cytometry data.

### Suppl. Figure 5

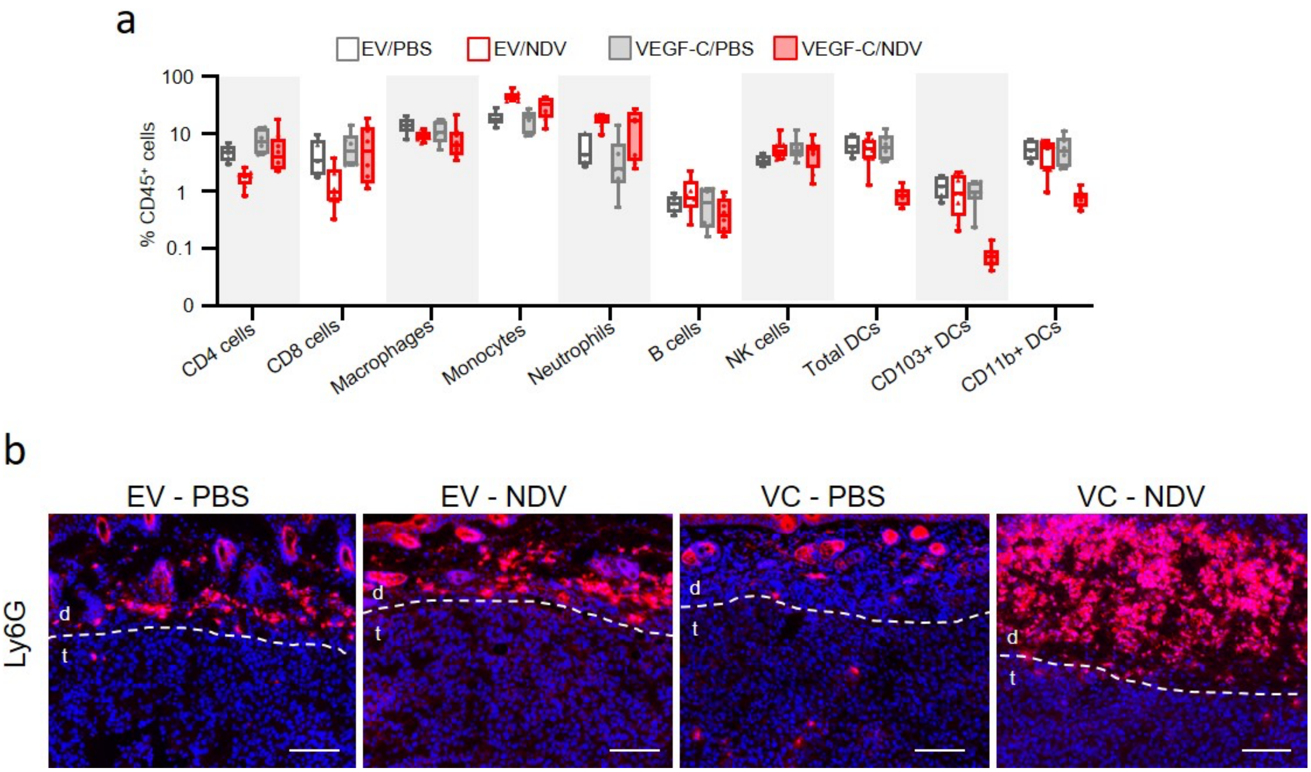


**Suppl. Fig 5. Immunophenotyping of NDV-treated B16/EV and B16/VEGF-C tumors** (a) Frequencies of distinct myeloid and T cell subtypes across treatments based on spectral flow cytometry data analyzed by manual gating. Frequences are shown as a percentage of total CD45 cells. Box-and-whiskers plots show individual values with mean ± SEM, and whiskers spanning the full range. EV/PBS, n=5; EV/NDV, n=8; VEGF-C/PBS, n=6; VEGF-C/NDV, n = 8. Statistical analysis with one-way ANOVA multiple comparisons Fisher LSD test. Refer to raw data for p-values. Outliers were removed based on ROUT analysis. Q=0.5%. (b) Immunofluorescent staining of tumors and adjacent skin for Ly6G+ neutrophils as indicated. d= dermis, t=tumor. Scale bar 100μm.

### Suppl. Figure 6

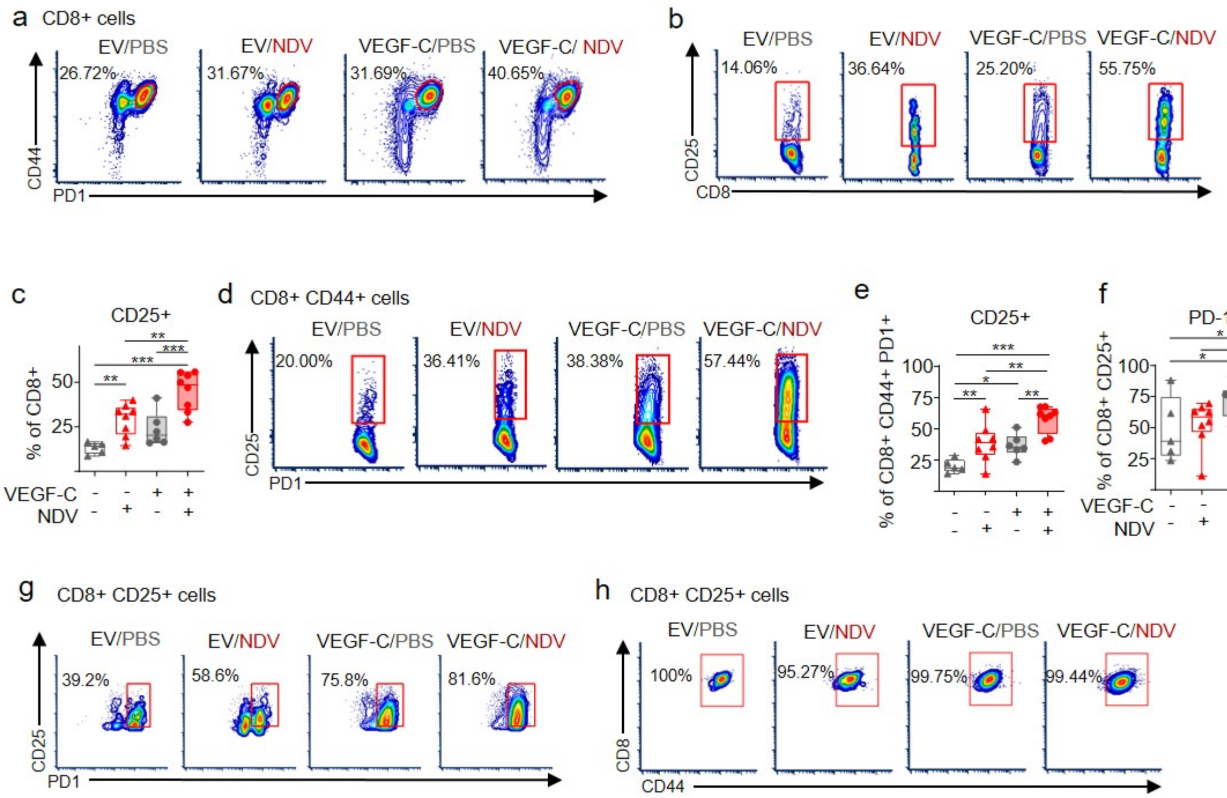


**Suppl. Fig.6. Flow cytometry analysis of $CD8^+$ T cells in tumors by manual gating.** (a, b) Contour plots of $CD44^+PD1^+$ (a) and $CD25^+PD1^+$ (b) effector $CD8^+$ T cells in B16F10 tumors across treatments as indicated. (c) Frequencies of $CD8^+CD25^+$ T cells. (d) Contour plots of $CD44^+CD25^+$ $PD1^+$ effector $CD8^+$ T cells. (e-h) Frequencies of $CD44^+CD25^+$ $PD1^+$ (e) and $CD25^+$ $PD1^+$ (f) effector $CD8^+$ T cells across treatments and corresponding contour plots (g, h). Box-and-whiskers plots show individual values with mean ± SEM, and whiskers spanning the full range. EV/PBS, n=5; EV/NDV, n=8; VEGF-C/PBS, n=6; VEGF-C/NDV, n = 8. Statistical analysis with one-way ANOVA multiple comparisons Fisher LSD test. . *$p < 0.05$, **$p < 0.01$, ***$p < 0.001$, not indicated = not significant.

#### Suppl. Figure 7

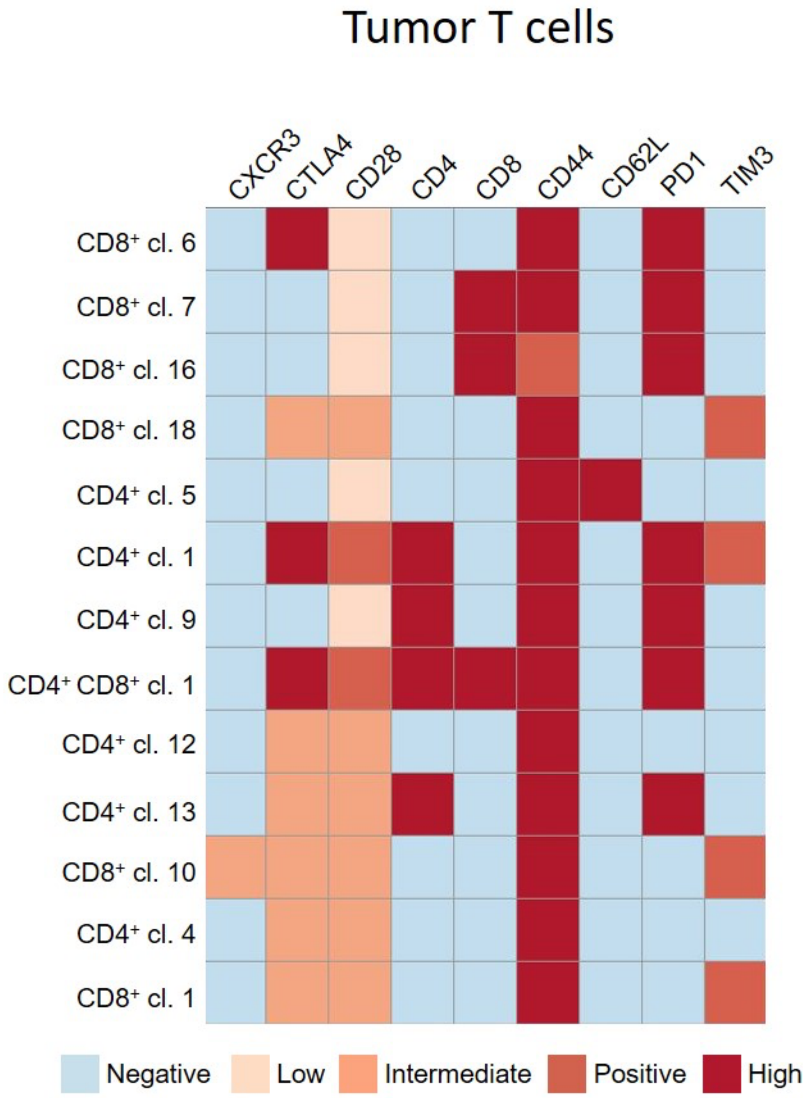


**Suppl. Fig 7. Heatmap showing T cell clusters and the corresponding cell surface markers.** Heatmap showing flow cytometry signal intensities for selected markers for the indicated cell populations from the tumor T-cell panel. Each square in the heatmap represents the median signal intensity of a marker across all cells within a specific cluster. The color scale reflects the level of marker expression, with signal intensities categorized into discrete levels based on marker-specific thresholds (see raw data file).

### Suppl. Figure 8

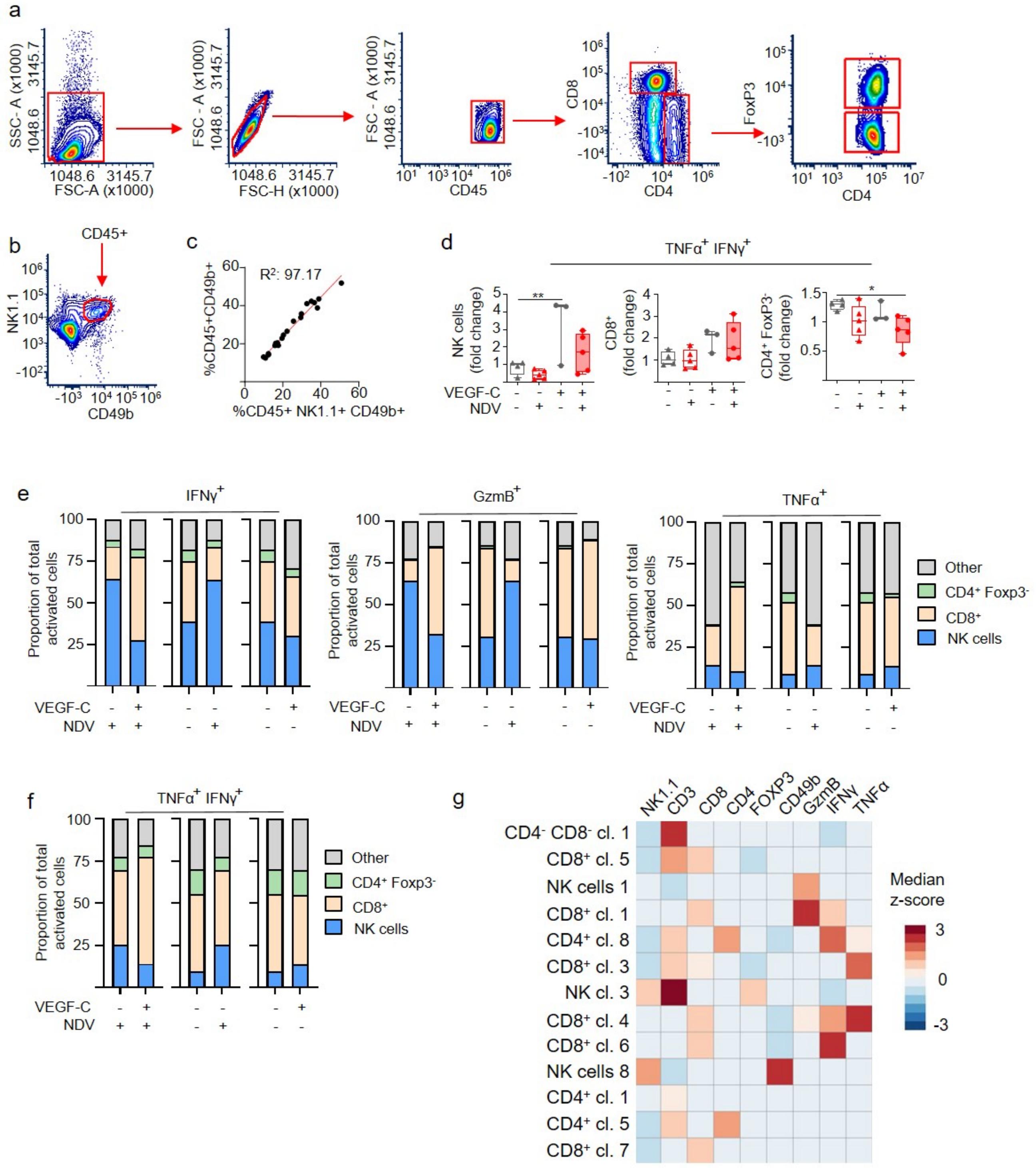

**Suppl. Fig. 8. Effects of VEGF-C and NDV on T cell and NK cell activation.** (a, b) Flow cytometry gating strategy for CD8$^+$, CD4$^+$ and NK cells. (c) Correlation analysis of NK cell markers CD49b$^+$ and NK1.1$^+$. (d) Increase or decrease (fold change) of NK cells, CD8$^+$ T cells, and CD4$^+$ conventional T cells, positive for both TNFα and IFNγ upon treatments as indicated. Box-and-whiskers plots show individual values with mean ± SEM, and whiskers spanning the full range. EV/PBS, n=4; EV/NDV, n=5; VEGF-C/PBS, n=3; VEGF-C/NDV, n=5. Statistical analysis with one-way ANOVA multiple comparisons Fisher LSD test. *$p < 0.05$, **$p < 0.01$, not indicated = not significant. (e, f) Proportion of T cells and NK cells positive for IFNγ, GzmB, or TNFα (e), or positive for both, TNFα and IFNγ (f), in B16/EV and B16/VEGF-C tumors across treatments. (g) Heatmap displaying median z-scores for the indicated populations, based on flow cytometry data using tumor intracellular panel.

### Suppl. Figure 9

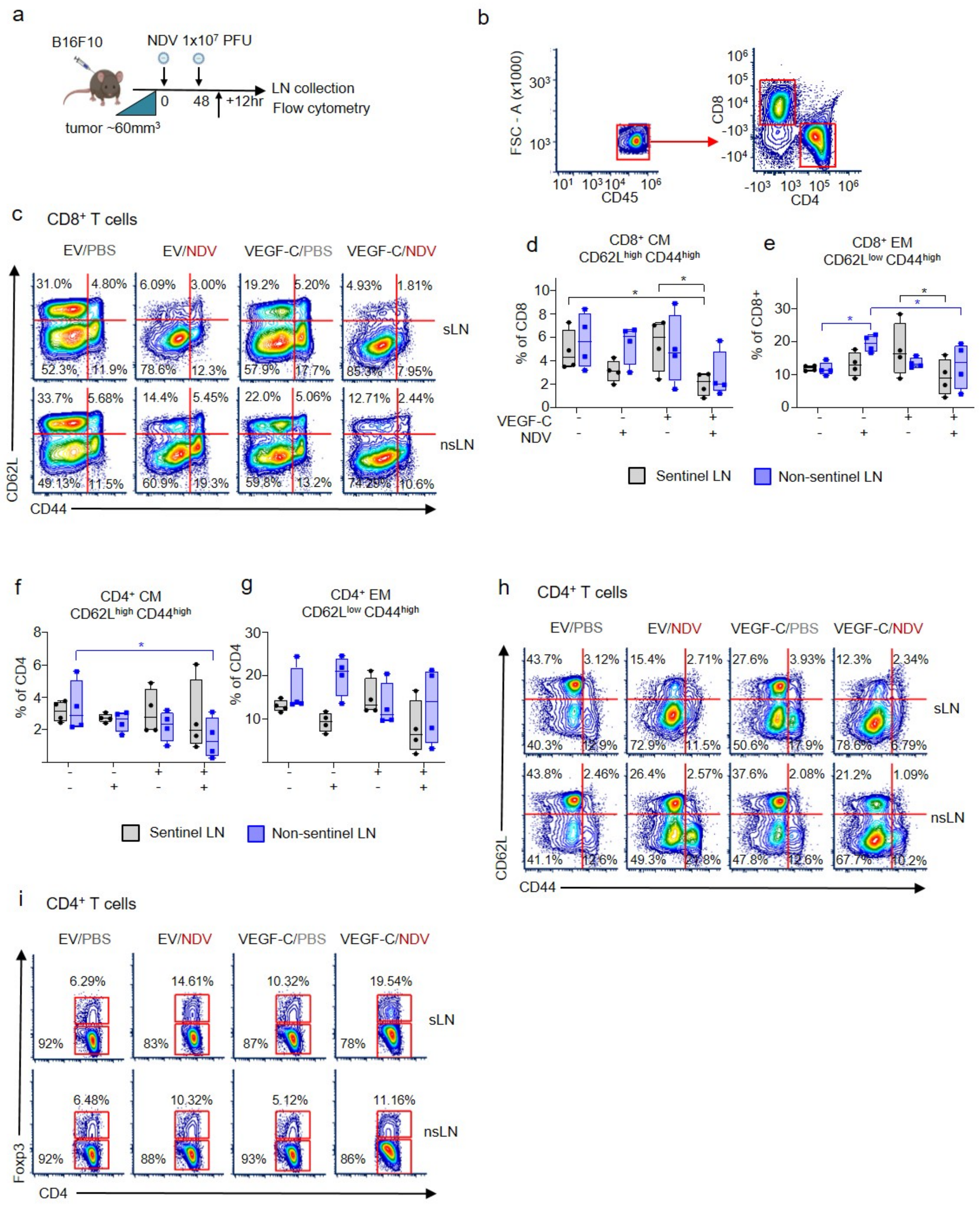

**Suppl. Fig 9. Characterization of T cell populations in sLNs and nsLNs of B16F10 tumors by flow cytometry** (a) Experimental design for sample collection: mice were injected with B16/EV or B16/VEGF-C cells and treated with NDV or PBS when tumors reached ~60mm$^3$. LNs were collected 12h after the second treatment. (b) Contour plots showing gating for CD8$^+$ and CD4$^+$ T cells in mouse lymph nodes. (c) Representative contour plots showing CD62L and CD44 expression on CD8+ T cells, as indicated. (d-g). Frequencies of effector memory (EM) and central memory (CM) CD8$^+$ and CD4$^+$ T cells in LNs. Box-and-whiskers plots show individual values with mean ± SEM, and whiskers spanning the full range.. Statistical analysis with one-way ANOVA multiple comparisons Fisher LSD test. n=4 for all groups. (h) Representative contour plots showing CD62L and CD44 expression on CD4$^+$ T cells, as indicated. (i) Representative contour plots for Tregs (Foxp3$^+$), and conventional (Foxp3$^-$) CD4$^+$ T cells upon treatments as indicated. *$p < 0.05$, **$p < 0.01$, not indicated = not significant.
